# A blueprint for high affinity SARS-CoV-2 Mpro inhibitors from activity-based compound library screening guided by analysis of protein dynamics

**DOI:** 10.1101/2020.12.14.422634

**Authors:** Jonas Gossen, Simone Albani, Anton Hanke, Benjamin P. Joseph, Cathrine Bergh, Maria Kuzikov, Elisa Costanzi, Candida Manelfi, Paola Storici, Philip Gribbon, Andrea R. Beccari, Carmine Talarico, Francesca Spyrakis, Erik Lindahl, Andrea Zaliani, Paolo Carloni, Rebecca C. Wade, Francesco Musiani, Daria B. Kokh, Giulia Rossetti

## Abstract

The SARS-CoV-2 coronavirus outbreak continues to spread at a rapid rate worldwide. The main protease (Mpro) is an attractive target for anti-COVID-19 agents. Unfortunately, unexpected difficulties have been encountered in the design of specific inhibitors. Here, by analyzing an ensemble of ~30,000 SARS-CoV-2 Mpro conformations from crystallographic studies and molecular simulations, we show that small structural variations in the binding site dramatically impact ligand binding properties. Hence, traditional druggability indices fail to adequately discriminate between highly and poorly druggable conformations of the binding site. By performing ~200 virtual screenings of compound libraries on selected protein structures, we redefine the protein’s druggability as the consensus chemical space arising from the multiple conformations of the binding site formed upon ligand binding. This procedure revealed a unique SARS-CoV-2 Mpro blueprint that led to a definition of a specific structure-based pharmacophore. The latter explains the poor transferability of potent SARS-CoV Mpro inhibitors to SARS-CoV-2 Mpro, despite the identical sequences of the active sites. Importantly, application of the pharmacophore predicted novel high affinity inhibitors of SARS-CoV-2 Mpro, that were validated by in vitro assays performed here and by a newly solved X-ray crystal structure. These results provide a strong basis for effective rational drug design campaigns against SARS-CoV-2 Mpro and a new computational approach to screen protein targets with malleable binding sites.

## INTRODUCTION

In December 2019, a new coronavirus (CoV), belonging to the clade b of the Betacoronavirus viral genus, caused an outbreak of pulmonary disease in the Hubei province in China.^1, 2^ In the first months of 2020, the new pandemic spread globally and it is still continuing. The virus shares more than 80% of its genome with that of the SARS coronavirus discovered in 2002 (SARS-CoV).^1, 2^ Hence it has been named severe acute respiratory syndrome-coronavirus 2 (SARS-CoV-2) by the International Committee on Taxonomy of Viruses.

The most promising tools for the cessation of the epidemic spread of COVID-19 are vaccines.^3^ The majority of them could trigger an immunogenic response using inactivated viral vectors or RNA sequences encoding SARS-CoV-2 spike glycoprotein.^4^ Unfortunately, the SARS-CoV-2 spike protein uses conformational masking and glycan shielding to frustrate the immune response, possibly hindering vaccine effectiveness.^5^

Interfering with viral replication is an alternative and promising strategy of treatment. In this context, the chymotrypsin-like proteinase (often referred to as the main protease, Mpro hereafter) is an excellent pharmaceutical target.^6, 7^ It does not depend on host immunogenic responses and it is essential for generating the 16 non-structural proteins, critical to the formation of the replicase complex.

Mpros are highly conserved enzymes across CoVs.^8, 9^ SARS-CoV Mpro was already suggested as one of the main drug targets in the pandemic associated with that virus, about 15 years ago.^10–12^ Inhibitors of proteases (e.g. aspartyl protease) are also common drugs used in the clinic against other deadly viruses, e.g. HIV-1.^13^

The SARS-CoV Mpro active form is a homodimer (Figure 1A), with each monomer consisting of N-terminal, catalytic and C-terminal regions^14^ (Figure 1B). Mpros were shown to process polyproteins on diverse cleavage sites, using a cysteine/histidine catalytic dyad:^15^ the histidine (His41 in SARS-CoV-2) forms a hydrogen bond (Hbond) with a water molecule that, in turn, interacts with an aspartate (Asp187) and a histidine (His164) side chain. Asp187 is further stabilized through a salt-bridge with a nearby arginine (Arg40, Figure 1C). In this way, His41 can act as a base, extracting a proton from the catalytic cysteine (Cys145) sidechain and activating it for the nucleophilic attack that cuts the polypeptide.

**Figure 1.**
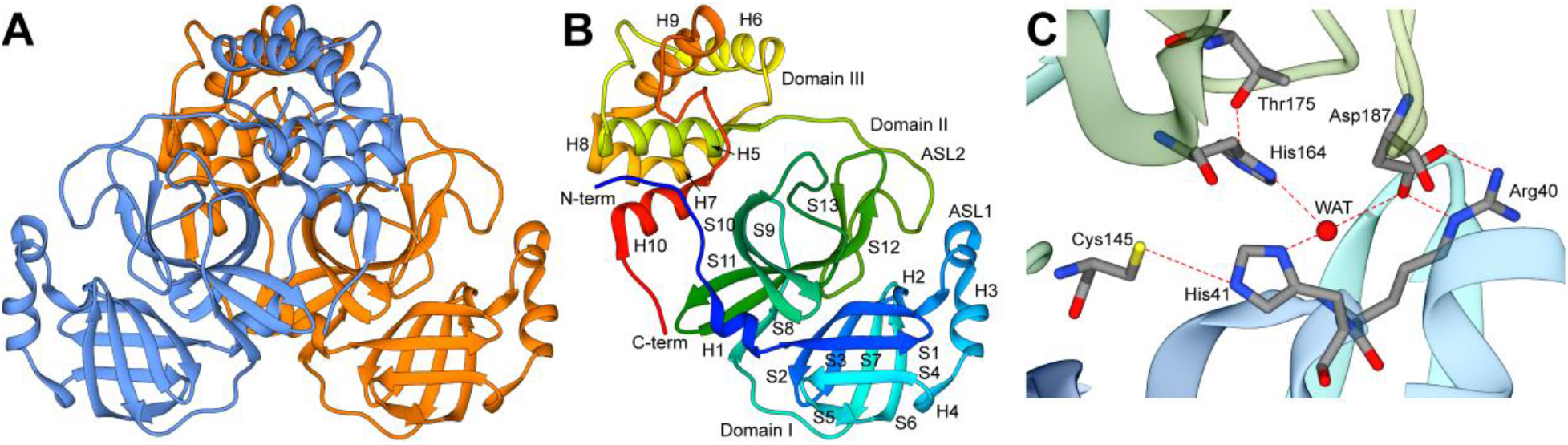
Structure of SARS-CoV-2 Mpro (PDB id: 6Y2E). (**A**) The enzyme is a homodimer.^17^ (**B**) Each monomer consists of three domains (I-III): The chymotrypsin-like and picornavirus 3C protease–like domains I and II (in blue and green respectively) form six-stranded antiparallel β-barrels, that harbor the substrate-binding site between them, including the ASL1 and ALS2 loops (residues 22-53 and 184-194, respectively). Domain III (in yellow-red) is a globular bundle formed by five helices and it is involved in the dimerization of the protein. (**C**) Close-up of the active site and of the Hbond network. Atoms are in stick representation colored according to atom type, while Hbonds are indicated using dashed lines.

SARS-CoV-2 Mpro shares 96% sequence identity with Mpro from SARS-CoV (Section S1, Figure S1A). Twelve residues differ between both Mpros and only one, namely, Ser46 in SARS-CoV-2 (Ala46 in SARS-CoV), is located at the mouth of the active site cavity (Figure S1B). The binding sites share 100% of sequence identity (Figure S1A). Thus, exploiting the known libraries of SARS-CoV Mpro inhibitors has been a strategy followed by many research groups. Unfortunately, most SARS-CoV Mpro inhibitors with good (nM) activity against SARS-CoV Mpro in vitro and in cell-based assays, exhibited limited (sub-μM) potency against the protein from SARS-CoV-2 in enzymatic assays, and low-μM IC50 values (4-5 μM) in cell-based assays.^16, 17^

A similar scenario has emerged from the virtual screening (VS) of Mpro inhibitors towards SARS-CoV-2. Indeed, none of these strategies: (i) repurposing of SARS-CoV drugs for SARS-CoV-2,^18, 19^ (ii) Deep Docking trained on SARS-CoV Mpro inhibitors,^20^ (iii) libraries of the other SARS proteases,^21–23^ and (iv) clinically approved drugs for other SARS Mpros or other similar proteases,^24–29^ led to clinical advances. Because only 12 residues, far from the binding site, differ between SARS-CoV Mpro and SARS-CoV-2 Mpro, the mutation of distant residues can substantially contribute to the binding site plasticity and to the ligand binding through allosteric regulation.^30^ This is both disappointing and puzzling from a pharmaceutical perspective. Recently, however, it has been shown that the dipeptide prodrug GC376, and its parent GC373 inhibit the two proteases with IC50 values in the nanomolar range.^31^ This suggests that, despite the intrinsic and significant differences between the two Mpros, common binding features against some classes of high-affinity ligands are retained.

Molecular dynamics (MD) simulations^18^ provided hints to address this riddle: they showed that the SARS-CoV Mpro active sites display major differences in both shape and size. In particular, while both Mpros reduce their accessible volume upon inhibitor binding by approximately 20%, the maximal volume of the *holo* SARS-CoV Mpro active site is over 50% larger than that of SARS-CoV-2. In addition, the accessibility of the binding hotspots (i.e. the key residues for substrate binding) and the flexibility of one of the two loops delimiting the binding pockets (ASL1 in Figure 1B) differs between the two Mpros.^18^ The simulations indicate that the binding sites of the two Mpros are dynamically diverse and that ligand binding can impact them differently.

Therefore, transferable binding features across Mpros, as well as unique ones for SARS-CoV-2 are difficult to predict: the exceptional flexibility and plasticity of the binding site is here coupled with large adjustments of the cavity shape in response to the binding of an inhibitor. This clearly emerges from an analysis of the SARS-CoV-2 Mpro binding pocket’s conformational changes (performed here) across the majority of the 196 X-ray crystal structures available in the Protein Data Bank up to September 30^th^ 2020.^18, 32–34^ This flexibility makes a rational drug-design approach extremely challenging:^18, 35^ the screening potential of Mpro conformational space is too large, too flexible, unpredictable, and the actual available binding space can differ significantly from ligand to ligand.^18, 32, 36^

It is therefore imperative to identify the relationship between SARS-CoV-2 conformational space, flexibility, druggability and ligand binding. Here, we analyzed the mentioned 196 X-ray crystal structures, along with about ~31,000 conformations extracted, not only from the longest (100 μs) MD simulation of SARS-CoV-2 MPro so far^37^, but also from binding site enhanced sampling simulations carried out here. Among these structures, we selected ~24,000 conformations for which we systematically performed druggability analyses on the binding sites. The top 110 druggable structures were selected for virtual screening of a sample library of ~12,600 ligands. The latter includes marketed drugs and compounds under development, the internal chemical libraries from Fraunhofer Institute and the Dompè pharma company, as well as the so-far known inhibitors of SARS-CoV Mpro from literature.

We redefine here the protein druggability in a new way exploiting the chemical space shaped by the different configurations of the binding site upon virtual screening. Specifically, we identified a consensus protein-ligand interaction fingerprint across the chemical space and the corresponding SARS-CoV-2 Mpro unique structure-based pharmacophore. Our pharmacophore was able to identify known nM-binders (IC50 ≤ 400 nM) of SARS-CoV-2 Mpro and to distinguish those from micromolar inhibitors. The latter was able to identify Myricetin and Benserazide as nM inhibitors, here experimentally validated by enzymatic activity binding assay. The predicted binding of Myricetin is also in striking agreement with the recently solved X-ray crystal structure by some of us. Our pharmacophore also provides a rationale for the variability in ligand affinity of currently known SARS-CoV ligands and for the general lack of transferability of SARS-CoV ligands to SARS-CoV-2, shedding light on the complexity, plasticity and druggability of SARS-CoV-2 Mpro.

## RESULTS

To understand how Mpro’s binding site conformation affects the druggability and the chemical space of selected binders, we start by considering ~30,000 Mpro conformations spanning different binding site arrangements and flexibility obtained from different sources (Table 1). These include:

i. All the X-ray crystal structures deposited to date of *apo* and *holo* SARS-CoV-2 Mpro. After analyzing all of them, we selected 43 structures for follow-up analysis. These exhibit a RMSD lower than 1 Angstrom with respect to the excluded ones. Therefore, they can be considered a good representative ensemble of the overall deposited SARS-CoV-2 X-ray crystal structures (“X-ray ensemble” hereafter, see Table S1 for details).
ii. Two sets of MD ensembles: (a) The “MD10000” ensemble, that is 10,000 frames, taken every 1 ns, of a 100 μs-long MD of *apo* Mpro from Shaw’s group^37^ and (b) the “MSM ensembles”, two collections of 30 and 40 representative conformations extracted from a three-state and a four-state Markov state model (MSM), respectively (see Section S2, Figure S2). The MSM analysis was performed on the 100 μs-long MD trajectory. These are included to identify conformations representing structural changes potentially related to binding.
iii. About 22,000 conformations obtained with two enhanced sampling methods implemented in the TRAPP web server:^38^ MD-based enhanced sampling by Langevin Rotamerically Induced Perturbation (LRIP)^39^ and constraint-based sampling by tConcoord^40^ as referred to as a “LRIP/tC ensembles” hereafter. These ensembles were derived using the X-ray crystal structures of the protein in complex with the inhibitor N3 (PDB ID 6LU7) and with another α-ketoamide inhibitor (PDB ID 6Y2G).

**Table 1.** Mpro conformational ensembles considered in this study.

| Mpro protein's conformational space |  |  |  |
| --- | --- | --- | --- |
| Name | Total number (N) of conformations analyzed | No. of binding sites analyzed (chain A and chain B) | No. of druggable binding sites screened (chain A and chain B) |
| X-ray ensemble | 196 | 86 | 84 |
| MSM ensemble (four-state model) /<br>MSM ensemble (three-state model) | 40 (four-state model)<br>30 (three-state model) | 80 (four-state model) | 20 (four-state model) |
| MD10000 | 10,000 | 2,000 | 16 + 40 |
| LRIP/tC ensembles | 21,994 | 21,994 | 16 + 40 |
| Ligands' conformational space |  |  |  |
| Name | Total numbers of molecules | Notes |  |
| Sample Library | 13534 | - |  |
| Active molecules | 193 | Fraction of the sample library for which an experimental IC50 measure against SARS-CoV is available |  |
| Crystallographic ligands | 37 | Available from the Protein Data Bank |  |
| Consensus molecules | 33 | Fraction of the sample library that is found in common across the top1% of the best-performing structures |  |
| Consensus active | 16 | Fraction of the active molecules found in the consensus molecules |  |
| SARS-CoV-2 Mpro binders | 41 | Available from literature |  |

### 1. Binding site features and druggability

Next, we calculated specific binding site parameters for the ensembles. These include the volume, the hydrophobicity/hydrophilicity and others (see Supplementary Section S3 for the complete list). We discuss only these two parameters, because they turn out to be key descriptors of the SARS-CoV-2 Mpro active site druggability (see below). The latter was evaluated with the “druggability” scores: SiteScore and Dscore, as derived from the SiteMap tool implemented in the Schrödinger suite 2019-4 (Schrödinger, LLC, New York, NY, 2019) and the CNN and LR (Convolution Neural Network and Linear Regression) druggability models^41^ as implemented in the TRAPP package.^42^ Although TRAPP and SiteMap use different approaches for computing the pocket characteristics (3D grid-based versus residue-based), the trends in the computed parameters are similar.

From this analysis, we conclude the following:

i. The binding site volumes computed with TRAPP for the *holo* X-ray crystal structures are distributed over a slightly larger and more variable range of values than that for the *apo* X-ray crystal structures (Figure 2A, Figure S3). The distribution of volumes is higher in the MD and enhanced sampling simulations: for instance, in the MD10000 ensemble, the volume of the *apo* protein has a variation of as much as 48% (Figure 2A) with respect to the average value. Similar trends were observed for volumes computed with SiteMap (see Table S2). This difference in volume distributions between X-ray crystal structures and MD snapshots could be caused by crystal packing. We therefore calculated the minimum distance between two crystallographic symmetry images, as well as the minimum distance between the binding site residues and the nearest image (see Section S3.2). Depending on the space group, the minimum distance between a non-hydrogen atom in the binding site and an image atom turns out to span between ~2.4 Å and ~9.3 Å (Table S3, Figure S5). Therefore, crystal packing might constrain the binding site of some of the crystal structures to more compact conformations. During the MD simulations, on the other hand, the large range and variability in binding site volume are associated with conformational changes of loops ASL1 (res. 22-53) and ASL2 (res. 184-194) (Figure 2B). It can be seen that ASL1 is more flexible than ASL2, from the MSM analysis (Section S2) and by calculation of the residue occurrence in the binding site (Section S3, Figure S3). Volume variability also results from the transient participation (with a frequency of ~25%) of the N- and C-terminal tails of the adjacent subunit in the binding site (Figure 2C). During the MD simulation, these two termini move in the proximity of the pocket. For the seven *apo* X-ray crystal structures, where the termini are resolved (Table S1), the C-terminus is never close to the binding sites, whereas the N-terminus is always so (see Figure 2D). In the *holo* crystal structures, the N- and C-terminal tails of the adjacent subunit can both be found close to the binding site (see Figure 2D). A similar scenario is observed for the *holo* structures in the LRIP/tC ensemble, where both terminals are present in the binding pocket even more often (in about 40% of simulated structures, Figure S3).
ii. The binding site hydrophobicity is here estimated in terms of hydrophobicity distribution across the different conformational ensembles, calculated with TRAPP. The distribution of the MD ensemble (based on the *apo* structure) can be fitted with a Gaussian centered around a hydrophobicity ~55 on the TRAPP scale^41^ (Figure S6A). Similarly, the *apo* X-ray crystal structures could be fitted with a Gaussian that is shifted to a peak at a higher hydrophobicity of ~65 (Figure S6A). The distribution of the *holo* X-ray crystal structures exhibits three peaks: one at lower hydrophobicity (~42-55), one at the *apo* X-ray crystal structures hydrophobicity (~60-70), and one at high hydrophobicity (~80-90) (Figure S6A). The high hydrophobicity conformations include complexes in which large ligands, with a molecular mass of 400 Da or greater, are covalently bound to Cys145 (PDB IDs 6LU7, 7BUY, and 7C8R). The hydrophobicity distribution from tConcoord can be fit with a more spread Gaussian spanning from low (~20) to high hydrophobicity values. The hydrophobicity distribution of the LRIP conformations is instead bivariate and overlaps with that of the MD ensemble. The observed trends for hydrophobicity computed with SiteMap for the crystal structure and MSM ensembles are similar, see Section S3.

**Figure 2.**
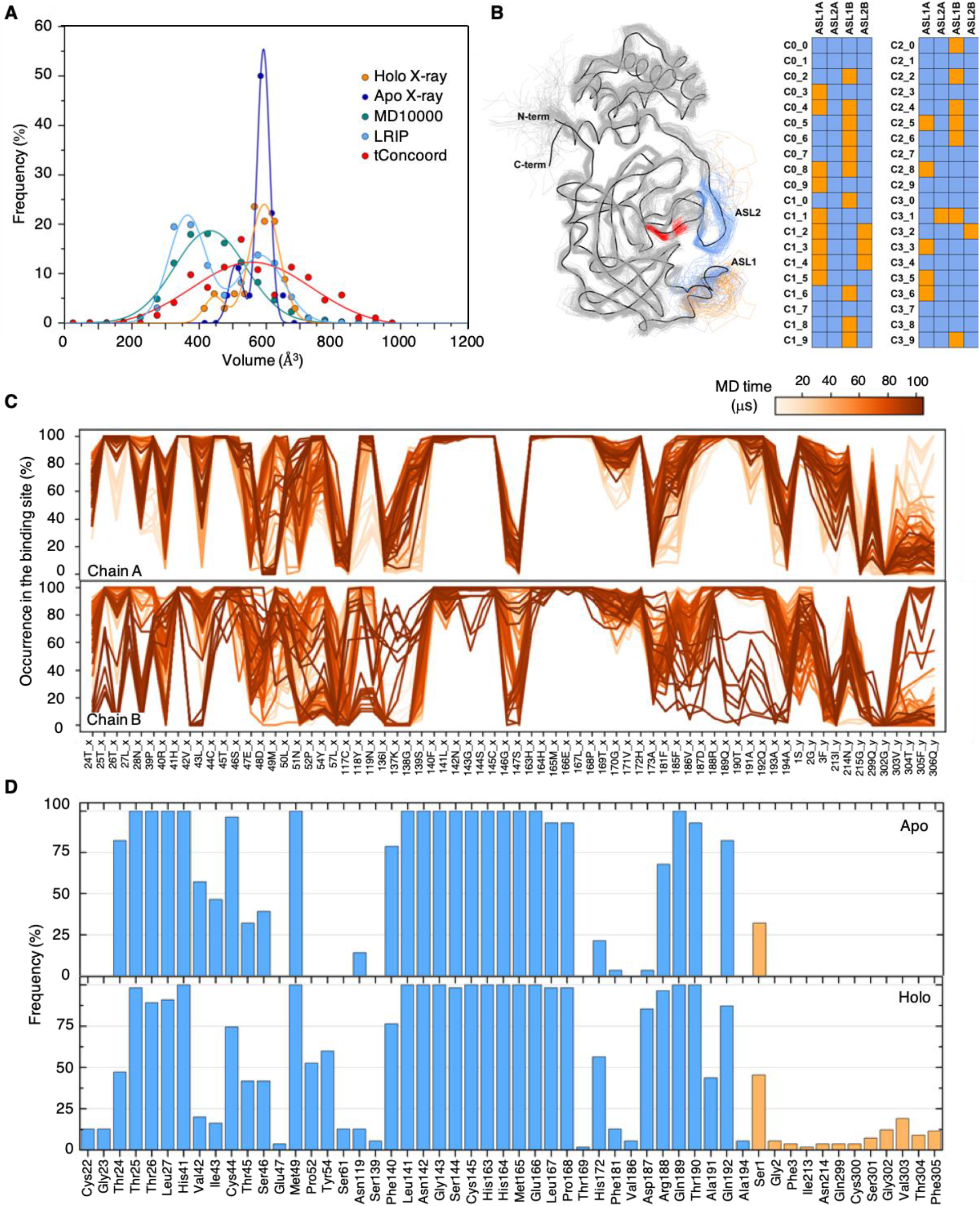
Binding site analysis of computational ensembles. (**A**) Binding site volume distributions from TRAPP for X-ray structures and MD/LRIP/tC ensembles. MD10000 contains conformations very similar to the frames extracted by the MSM analyses on the same trajectories, therefore these are not shown here. Binding site volume distributions from SiteMap are in Figure S3. (**B**) Analysis of ASL1 and ASL2 conformations (in chain A and B) in MSM (four-state model). Orange and blue squares refer to open and close conformations, respectively. The three-state model clusters are reported in Figure S3. (**C**) Binding site residues in the 100 μs-long MD trajectory: average occurrence in snapshots at 1 μs intervals for chains A and B separately. Unlike chain A, extensive conformational changes are observed in the loop formed by residues 181-194 of chain B (see Figure S4). (**D**) Average occurrence of the binding site residues in the set of apo and *holo* X-ray structures (Table S1). Residues from the same chain are shown in blue, while residues from the adjacent chain are colored in orange. The percentage of each residue was calculated considering the number of structures for which that residue was resolved.

#### 1.1. Druggability

Here, we analyze the druggability as defined by the CNN and LR scores from TRAPP^41^ along with DScore/ SiteScore from SiteMap.^43, 44^ All pockets in the crystallographic structures (except one, PDB ID 6WTK) are scored as druggable: their druggability indices are above the scores’ thresholds for druggability (0.9, 0.9, 0.5, 0.8 for SiteMap, DScore, LR and CNN models of TRAPP, respectively, Figure S6. These thresholds were taken from Halgren *et al.*^44^). There is not, however, any notable correlation between the druggability scores from the SiteScore/DScore and TRAPP methods. This is expected because the observed variations in the druggability index is within the method prediction uncertainty. Despite the slightly lower druggability indices in SiteScore and TRAPP-LR for the *apo* X-ray crystal structures compared to the *holo* crystal structures, they are still predicted to be druggable within the uncertainty of the methods. In contrast, about 50% of the simulated structures (MD, LRIP, and tConcoord) were predicted not to be druggable (Figure S6).

The druggability scores of the simulated, and, more, of the X-ray crystal structures correlate with binding site hydrophobicity (Figure S6, Section S3.3, and Table S4, S5). The correlation with other binding site features is much smaller (see Table S4, S5). We conclude that, as expected, the more hydrophobic the pocket is, the more druggable it is.

### 2. Virtual screening

We defined a sample library of a total of 13,535 compounds (Table 1, Figure S7). The library included commercialized drugs and compounds under development, the internal chemical library from Dompè pharma company and compounds from the Fraunhofer Institute BROAD Repurposing Library, as well as known inhibitors of SARS-CoV-1 Mpro. In particular the library included a set consisting of 180 compounds with pIC50 against SARS-CoV Mpro greater than 6 reported in the literature (active molecules, hereafter).^12, 16, 45–67^ Our sample library is chemically very diverse as compared to the crystallographic ligands in complex with SARS-CoV-2 Mpro and the active molecules (see Figure S7). A more detailed chemoinformatics analysis is reported in Section S4.1-2.

We selected SARS-CoV-2 Mpro conformations with scores above the druggability thresholds. These include all the X-ray crystal structures except 6WTK (42 structures), 10 MSM ensemble conformations (MSM selection), and 8 representatives of the top 10% scoring conformations from the MD10000 and LRIP/tC ensembles (see Table 1, Section S4.3, Table S6, Figure S8-9).

The ligands were screened against the conformations using OpenEye FRED^68^ and Schrödinger Suite Glide Version 85012.^69, 70^ We discuss here the results obtained with FRED. Those obtained with Glide present similar trends and are reported in the Supporting Information (Section S4.4 and Figure S10). Also, we report here only the calculation results obtained with the MD10000/LRIP/tC and X-Ray selections. The data obtained with the MSM selection are reported in the SI (Section S4.5, Figure S11).

The quality of the virtual screenings was evaluated in terms of: (i) Enrichment Factor (EF) defined as EF(1%) = (#active molecules in the top 1% / #molecules in the top 1%) / (active molecules in the whole set). (ii) Receiver Operating Characteristic (ROC) curves, used to evaluate the true positive rate and the area under the curve (AUC).

The structures from the MD10000/LRIP/tC and MSM selections, along with the *apo* X-ray structures, exhibited a poor EF (below 5%), despite being identified as druggable by all the druggability prediction methods here implemented (Figure 3). The poorer performance of the *apo* X-ray crystal structures was expected since they exhibited overall lower druggability scores with respect to the *holo* structures (Figure S6F). This was not the case with the selected structures from the MD10000 and LRIP/tC ensembles (see Section S4.3), where only the 10% of the structures with highest druggability scores were used for screening. This suggests that the druggability prediction methods are not sensitive enough to distinguish between high- and low-EF conformations for the chemical space considered (see also discussion below). On the other hand, for the *holo* X-ray crystal structures, both “well-performing” (EF > 15%) and “poorly-performing” (EF < 5%) conformations were identified (Figure 3).

**Figure 3.**
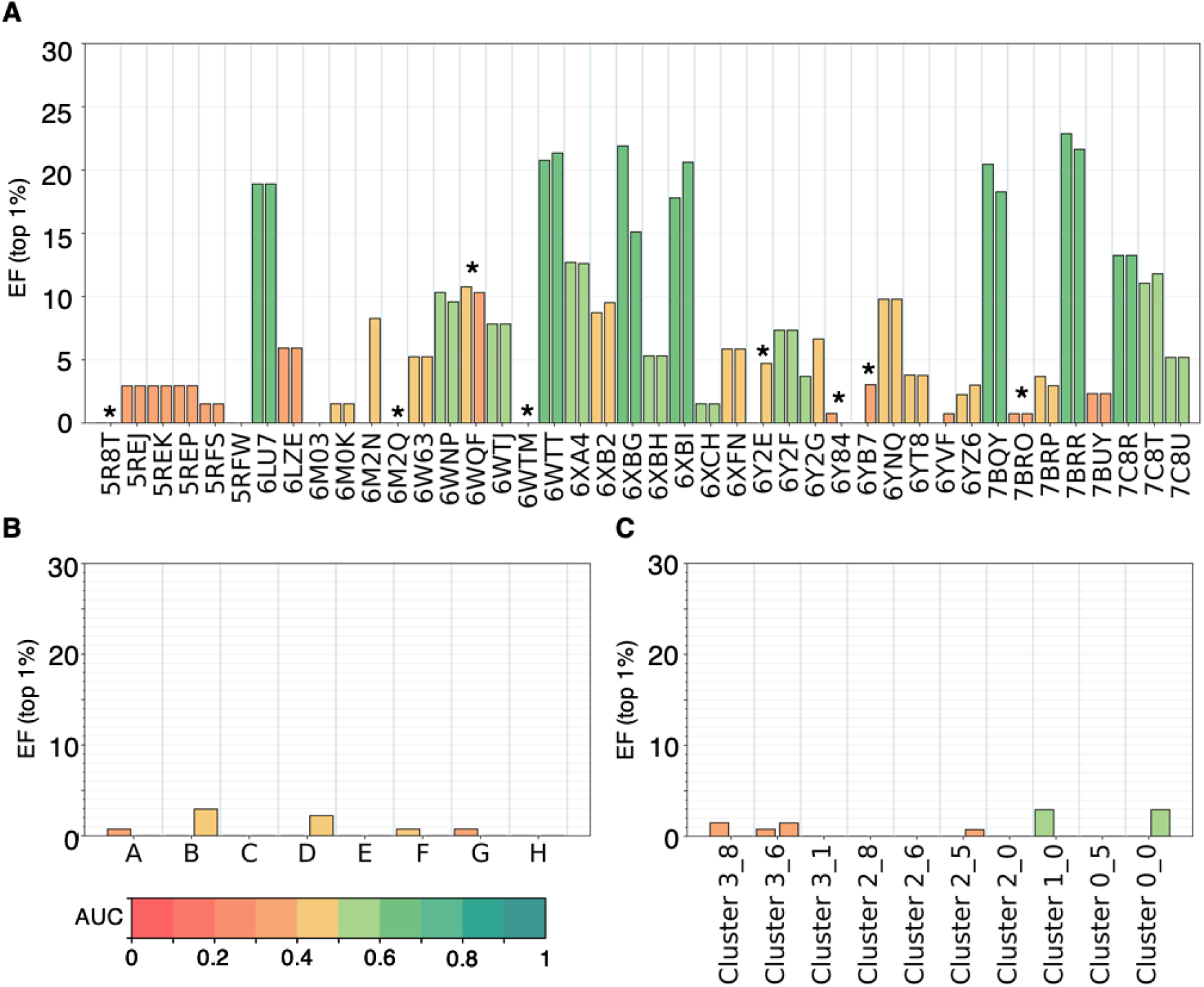
Virtual screening performance evaluation. Top 1% EF from the FRED virtual screenings performed on the binding sites in chains A and B in the X-ray crystal structures (**A**), in the MD10000 and LRIP/tC selection (Section S3.4) (**B**) and in the MSM selection (**C**), i.e. 10 structures with druggability index > than 0.9 as extracted from the four-state model MSM ensemble. The bars are colored from red to dark green according to the value of the Area Under Curve (AUC) in the ROC curves (color scale on the bottom left corner). The “*” symbol in panel (**A**) highlights the apo structures. Glide results are shown in Figure S10.

Next, we determined which of these ensembles exhibits a Protein-Ligand Interaction Fingerprint (PLIF) comparable to the one established by the ligands co-crystallized with SARS-Cov2 Mpro. In the PLIF of the latter (Figure 4A), we observe an overall predominance of Hbond interactions over hydrophobic ones, with Cys145 (catalytic dyad), Gly143, Ser144, His163 and Glu166 as the most attractive residues to form Hbond interactions (comparable occurrence) with the ligands. The only exception is represented by His41 (catalytic dyad), which is similarly involved in Hbonds and hydrophobic interactions.

**Figure 4.**
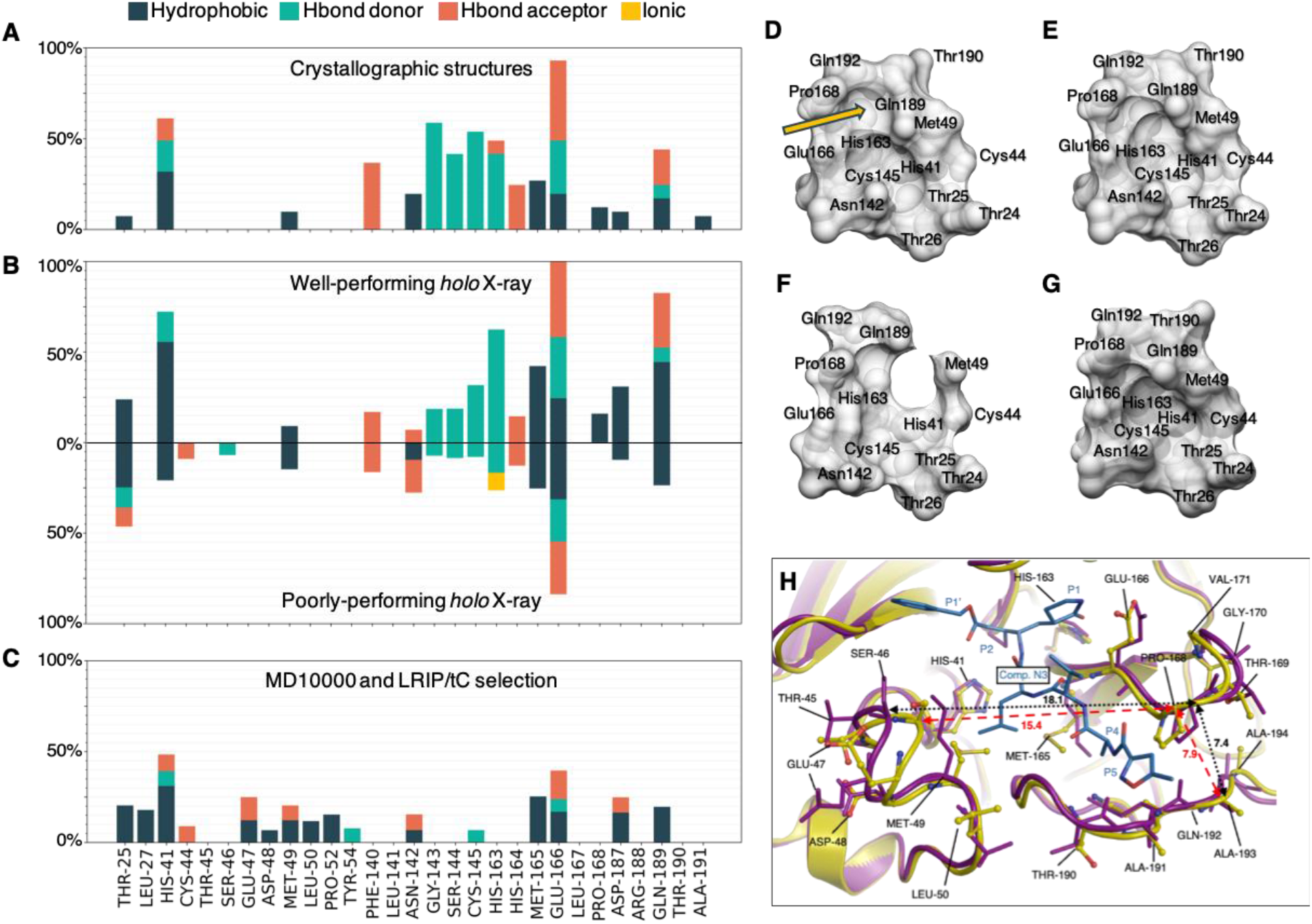
Virtual screening pose analysis of well- and poorly-performing receptor conformations. Average PLIF of (**A**) the crystal structures, (**B**) the top 1% of molecules from virtual screenings on well- and poorly-performing X-ray structures and (**C**) the MD10000 and LRIP/tC selection. The PLIF of the top 1% of the MSM selection is reported in FigureS11. The occurrence of interactions between the ligands and the well- and poorly-performing structures is plotted on the upper and lower half-plane, respectively. All bar plots were normalized with respect to the highest found occurrence (interactions with Glu166 in upper panel **B**). Binding site shape averaged over the well-performing (**D**), poorly-performing (**E**), the MD10000/LRIP/tC selection (**F**) and the apo (**G**) structures (**H**). Superposition of a well- and poorly-performing crystal structures of SARS-CoV-2 Mpro (6LU7 and 5REK, well- and poorly-performing, respectively); ribbons are in purple and in yellow, respectively, while the N3 ligand in blue carbon stick representation. The Ser46-Pro168 and Ala193-Pro168 Cα distances are highlighted. This latter panel was based on the scheme published in Kneller *et al.*^30^

The PLIF of the well-performing conformations (Figure 4B, upper panel) matches the one of the crystallographic complexes well (Figure 4A). Namely, the same hot-spot (i.e. preferential residues for ligand binding) residues emerge: the ligands form Hbonds with Cys145 (catalytic dyad), Gly143, Ser144, Gly166, and His163, as well as hydrophobic interactions with His41 (catalytic dyad). The main difference between the two PLIFs is in the lower occurrence of Hbonds involving Gly143, Ser144 and Cys145 (catalytic dyad) than in the crystallographic complexes, and in the higher occurrence of hydrophobic contacts with Thr25 and His41 (the second residue in the catalytic dyad). This change in the surrounding of the reactive cysteine, Cys145 (Thr25, Gly143, Ser144) might be due to the presence of several covalent ligands in the crystal structures. Covalent binding might locally alter the PLIF, and this effect is not considered in the virtual screening.

In all the other selections (i.e. poorly-performing X-ray structures Figure 4B lower panel, MD10000 and LRIP/tC selection Figure 4C, and MSM selection Figure S11), the key Hbonds above discussed have an occurrence that is markedly lower than non-specific hydrophobic interactions. Also the latter interactions substantially decrease, including those with His41 (catalytic dyad). Moreover, in the MSM, MD10000 and LRIP/tC selections, ligands interact with almost all residues of the binding site (Figure 4C and Figure S11), but with an occurrence below 25% (for each interaction type) and with a strong predominance of hydrophobic interactions versus HBonds. This points to a rather non-specific binding of the screened molecules in the MD/MSM-selected structures.

Summarizing, we found that our evaluation of the virtual screening procedure correctly identifies the conformations able to provide the most similar PLIF to the known crystallized ligands of SARS-CoV2 Mpro.

To rationalize why the binding site structural determinants cause such dramatic differences in the PLIF of crystal structures (i.e. poorly/well-performing), MSM, MD10000 and LRIP/tC selections, we compared the average binding site shapes for the different selections. We found that the residues in the MSM, MD10000 and LRIP /tC selections are distributed over a larger volume than in the crystal structures (Figure 4D-F); therefore, the spatial location of the hotspots (i.e. HBond donors/acceptors, hydrophobic patches, charges) is significantly different. MSM, MD10000 and LRIP/tC selections were composed of structures with high druggability scores (see Section S3.3), suggesting that the druggability scores are, in this case, unable to identify the binding site features responsible for good performances (defined here as EF and AUC, see above) in virtual screening. Even re-selecting the conformations from the MD-ensembles using the similarity with respect to well-performing structures as criterion (i.e. Root Mean Square Deviation, RMSD), did not provide a satisfying performance (Section S5, Figure S12), indicating that key features for obtaining high enrichment factors in the virtual screening, i.e. the precise placement of interacting residues, are missing.

Let us now compare the average binding site shape of the well-versus the poorly-performing crystal structures: two adjacent cavities can be observed on the well-performing structures (Figure 4D): one including Glu166, Pro168, Gln189, Thr190 and Gln192, that is missing in the poorly-performing structures (Figure 4E); the other including Thr25 Leu27, His41, Asn142, Cys145, Met165, Glu166 and Gln189 that is also present in the poorly-performing structure, but wider and less deep than in well-performing ones. Therefore, interactions with Pro168 only appear in the well-performing structures, and the occurrence of Gln189 is much higher in well-performing than in poorly-performing ones. Notably, the binding site shape of the poorly-performing structures shares strong similarities with that of the *apo* structures (Figure 4G).

This difference in binding site shape is a consequence of the fact that in the well-performing structures, the small helix near P2 group (residues 46–50) and the β-hairpin loop near P3–P4 substituents (residues 166–170) shift about 2.7 Å apart with respect to the poorly-performing ones, whereas the P5 loop (residues 190–194) moves closer to the P3–P4 loop. Also, two methionines, Met49 and Met165, change their side-chain orientation impacting the side chain positions of the overall P2 loop, as well as the exposure of His163 (Figure 4H); the interaction with the latter significantly decreases in the poorly-performing structures, as already discussed above. These differences are similar to those observed between the well-performing and the *apo* structures (Figure S13).

Taken together, these results may explain why the druggability indices are unable to distinguish the binding site features linked to a good performance in virtual screening: very subtle variations of the conformations of the binding site residues induced upon binding, and therefore not present in the structural ensembles generated in the absence of a ligand, can lead to significant differences in the EF. These subtle variations are very challenging to discriminate in terms of druggability indices.

### 3. The active-pharmacophore and the SARS-CoV-2 blueprint on the chemical space

To identify the relevant ligand features that might relate to high affinity binding, we extract here the consensus chemical space, defined by the common ligands across the top 1% of the well-performing structures in the virtual screening. These are 32 molecules, 16 of which belong to the “active” molecules in our library (“consensus active” hereafter, Figure S14 and Table 1). We next calculated the corresponding pharmacophore (i.e. an ensemble of steric and electronic features that is necessary to ensure the optimal supramolecular interactions with a specific biologic target) and linearly combined it with the X-ray pharmacophore. This is done to consider possible additional features coming from covalent binding that cannot be covered by the virtual screening protocol. The consensus pharmacophore combined with the X-ray pharmacophore constitutes the “active-pharmacophore”, hereafter (see Methods, Figure 5).

**Figure 5.**
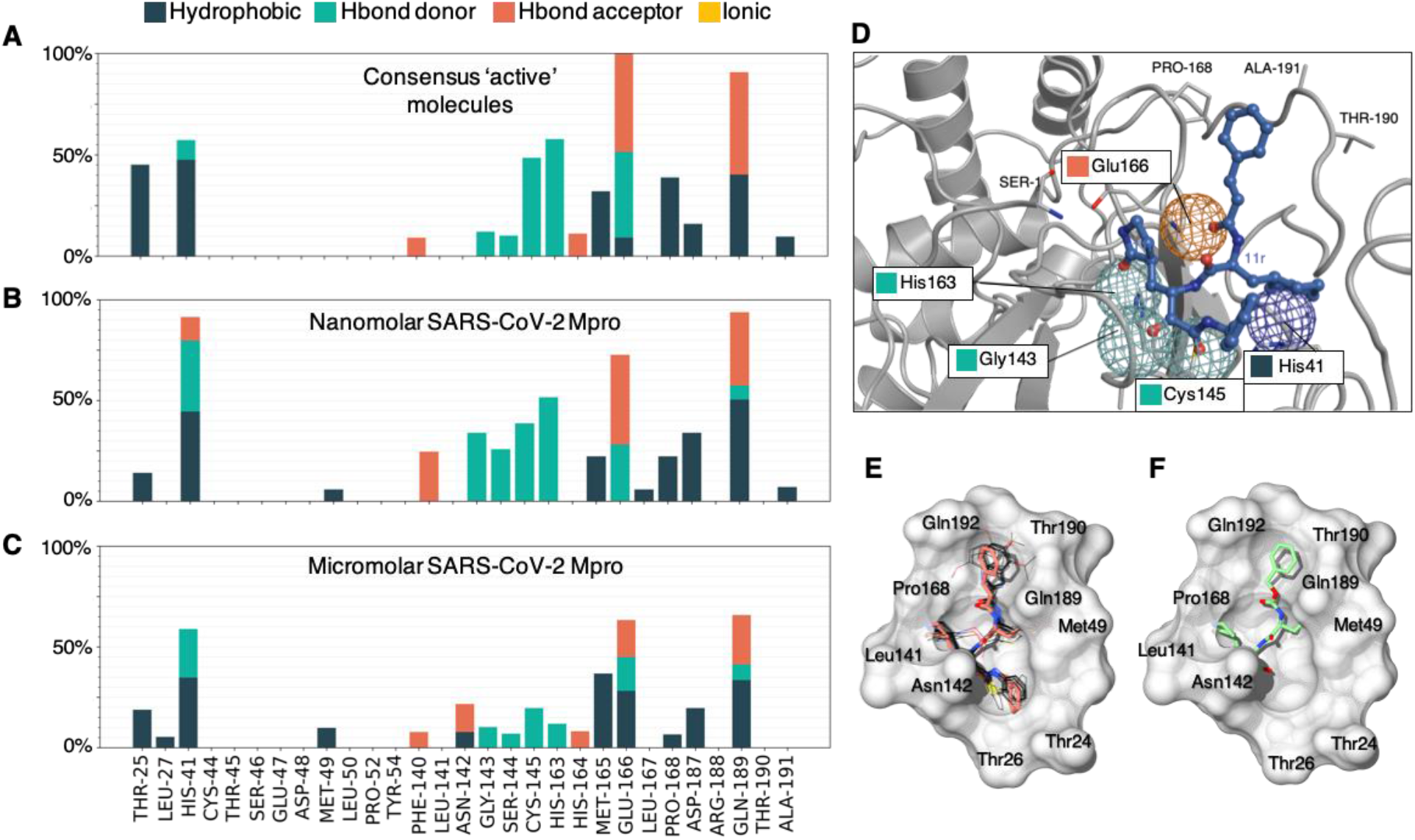
Virtual screening pose analysis of “consensus active” molecules. Average PLIF of the (**A**) 16 “consensus active” molecules, (**B**) 7 nanomolar (IC50 ≤ 400 nM) inhibitors and (**C**) 19 micromolar inhibitors docked onto the well-performing receptors. (**D**) Active-pharmacophore, where the 5 fundamental interactions (according to the selected cut-off, see methods) are displayed as spherical meshes. The docked pose of 11r, satisfying all the 5 interactions, is shown. (**E**) Docking poses of 16 “consensus active” molecules in SARS-CoV-2 Mpro (PDB ID 6LU7, chain A) binding site (black carbon representation). The 11r inhibitor pose is superimposed and highlighted in orange. (**F**) Docked pose of the inhibitor GC373.

Next, we tested the predictive power of our active-pharmacophore in discriminating the higher affinity binders across all the so-far known SARS-CoV-2 Mpro inhibitors: these are 46 molecules coming from the papers published until November 20^th^ 2020, which were not included in our sample library and that display a measured affinity spanning from 30 nM to 125 μM^17, 31, 71–77^). Some of these molecules were excluded by the docking software due to their excessive size or due to the presence of metals (e.g. candesartan cilexetil, Evans blue, phenylmercuric acetate).

For this purpose, we calculated the Dice coefficient, which measures the number of features in common between the molecule and the active-pharmacophore, relative to the average number of features present.^78^ When scoring the 46 known SARS-CoV-2 Mpro binders according to the Dice coefficient, the highest scored molecules were 11a, 11b, 11r, UAWJ246, UAWJ247, UAWJ248 and CG373 (see Figure S15), which are also the highest affinity (IC50 ≤ 400 nM) SARS-CoV-2 Mpro binders (nM-binders, hereafter). On the other hand, it is not possible to discriminate the sub-μM-binders of SARS-CoV-2 Mpro (400 nM < IC50 ≤ 1000 nM) from the μM ones (IC50 > 1000 nM) (see Figure S16). Of course, these results have to be taken with care given the fact we are comparing assay-dependent IC50 values coming from different labs. Also, several of these inhibitors are predicted to be covalent binders and, therefore, their IC50 values can be controversial (see the discussion offered in the Limitation paragraph). Therefore, in the next section, we analyzed the chemical space shaped by the well-performing conformations upon ligand binding, and offer a rationale for the predictive power of our active-pharmacophore in identifying nM-binders of SARS-CoV-2 Mpro.

#### 3.1. Rationalization of the active-pharmacophore and SARS-CoV to -CoV-2 Mpro ligands’ transferability

The PLIF of the consensus chemical space is dominated by the “consensus active” ones (the latter PLIF is almost identical to the former one, Figure S17A) and it shows a predominance of HBond interactions with His163, Glu166, Gly189 and Cys145 (Figure 5A), as well as hydrophobic interactions with Thr25, His41, Met165, Pro168 and Gln189. Accordingly, the same trends can be seen for the PLIF of the SARS-CoV-2 Mpro nM-binders (Figure 5B), the only differences being (i) the additional high occurrence of Gly143 and Ser144 Hbonds (as already observed in the PLIF of the X-Ray structures), and (ii) the lower occurrence of hydrophobic interactions with Thr25.

When comparing the binding poses of the “consensus active” molecules and the nM-binders of SARS-CoV-2 Mpro (Section S6), we found indeed that: the indole group (in 11a, 11b and 7 of the 16 molecules of the “consensus active” set) or the benzyl group (in GC373, 11r and 6 of the 16 molecules of the “consensus active” set), or the benzimidazole group (in 1 of the 16 of the “consensus active” set) is buried in the upper sub-cavity defined by residues Glu166, Pro168, Gln182, Gln189 and Thr190 (Figure 5E-F). Notably, this cavity was shrunk in the poorly-performing structures, further validating the quality of our model that correctly excluded the conformations potentially incompatible with nM-binders. The benzothiazole moiety of the “consensus active” is instead located in the lower part of the binding cavity defined by Thr24, Thr25 and Thr26. This benzothiazole moiety is absent in the SARS-CoV-2 Mpro nM-binders, possibly explaining the lower occurrence of hydrophobic interactions with Thr25.

This suggests that the binding to the lower part of the binding site (Thr24, Thr25, Thr26) is not a relevant feature for the nM affinity of SARS-CoV-2 ligands. In contrast, the high occurrence of Gly143 and Ser144 Hbonds appears to be a signature of nM-binders of SARS-CoV-2 Mpro, also found in the PLIF of the known X-ray ligands of SARS CoV-2 complexes. Notably, the formation of these two Hbonds appear to be significantly hampered in the “consensus active” set due to the presence of the above-mentioned benzothiazole moiety, that seems to compromise the juxtaposition of the Hbond acceptors of the ligands. Accordingly, none of the SARS-CoV-2 nM-binders, display benzothiazole or analogous bulky aromatic groups in such a position (Figure S14).

Analysis of all the 166 *holo* SARS-CoV-2 Mpro crystal structures also showed that the majority of their ligands do not have a benzothiazole or analogous bulky aromatic groups in the Thr24, Thr25, Thr26 subpocket (Figure S18). The two exceptions are PDB IDs 7JKV and 6XR3, where the ligands are covalently bound and the bulky group appear twisted in the cavity in a different position with respect the predicted the other X-rays and docking poses: this is possibly due to ligand-dependent conformational rearrangement upon covalent bond formation with Cys145. Concerning the μM binders known so-far, only very few of them are predicted to have a bulky group in such a position (Figure S16). In other words, our results suggest that the bottom part of the binding cavity in SARS-CoV-2 Mpro should only host small aromatic/hydrophobic moieties (or nothing at all) to facilitate the formation of Gly143 and Ser144 Hbonds, the latter being a signature of the currently known nM-binders to SARS-CoV-2 Mpro.

Recently, the X-ray structure of GC373 in complex with SARS-CoV-2 Mpro was solved (PDB ID: 7BRR, recently superseded by PDB ID 7D1M (released on October 28^th^ 2020). The ligand in the crystal structure appears in two different conformations, one resembling the predicted pose from us, where the benzyl group of GC373 is not buried in the upper sub-cavity defined by residues Glu166, Pro168, Gln182, Gln189 and Thr190; The other where this benzyl group it is exposed toward the solvent. Yet, when the crystallographic complex undergoes 500 ns of MD simulations, the pose where the aromatic ring is exposed toward the solvent rearranges as in the predicted docking pose (see Section S7 and Figure S19). Such results further validate our active-pharmacophore.

We next calculated the PLIFs of both the sub-μM (Figure S17B) and μM inhibitors (Figure 5C) of SARS-CoV-2 Mpro, which we were unable to discriminate with our active-pharmacophore. We found that indeed the two PLIFs are very similar and they both feature a drastic decrease of the occurrence of all the key HBonds found by the nM-binders of SARS-CoV-2 Mpro. The sub-μM and μM PLIFs turn out to resemble the PLIF of the “active” molecules that were not in our consensus group (i.e. nM-binders for SARS-CoV Mpro predicted to be low affinity binders for SARS-CoV-2 Mpro, Figure S20A, Section S8). Indeed, the “active” molecules excluded by our consensus present differences that may hamper a positive scoring and, possibly, a successful binding to the receptor, like an excessively long peptidic chain (i.e. Figure S20, molecules a and d), bulky substituents (i.e. Figure S20, molecules a and b) or a five membered ring rarely observed in Mpro inhibitors (i.e. Figure 20, molecule c).

This suggests that our active-pharmacophore is not only able to discriminate the nM-binders of SARS-CoV-2 Mpro (IC50 ≤ 400 nM) from the rest, but it also identifies key specific transferable and not-transferable binding features of nM SARS-CoV Mpro binders to SARS-CoV-2 Mpro ones. Taken together, these results suggest that our active-pharmacophore is a fair representation of the SARS-CoV-2 Mpro blueprint in the chemical space. Namely, it correctly represents a set of binding features compatible with the induced SARS-CoV-2 conformational space of the binding site. The latter is in part determined by the ligand upon binding and in part it depends on the residues differing in SARS-CoV-2 Mpro with respect to SARS-CoV Mpro, as also shown in Bzówka *et al.*^18^

### 4. Identification of nM-binders of SARS-CoV-2 Mpro

We considered a set of publicly available compounds within the E4C network,^79^ from the EU-OPENSCREEN Bioactive CompoundLibrary,^80^ coming from the PROBE MINER repository.^79, 81^ The set was re-scored based on the Dice similarity of their docked pose to our active-pharmacophore (see Methods). Benserazide (EOS100736) and Myricetin (EOS100814) compounds (see schemes in Figure 6A, B) were predicted as nM-binders of SARS-CoV-2 Mpro candidates. SARS-CoV-2 Mpro biochemical assays performed here established the accuracy of our predictions by measuring IC50 values as low as 140 nM and 220 nM, respectively (see Section S9, Figure S21).

**Figure 6.**
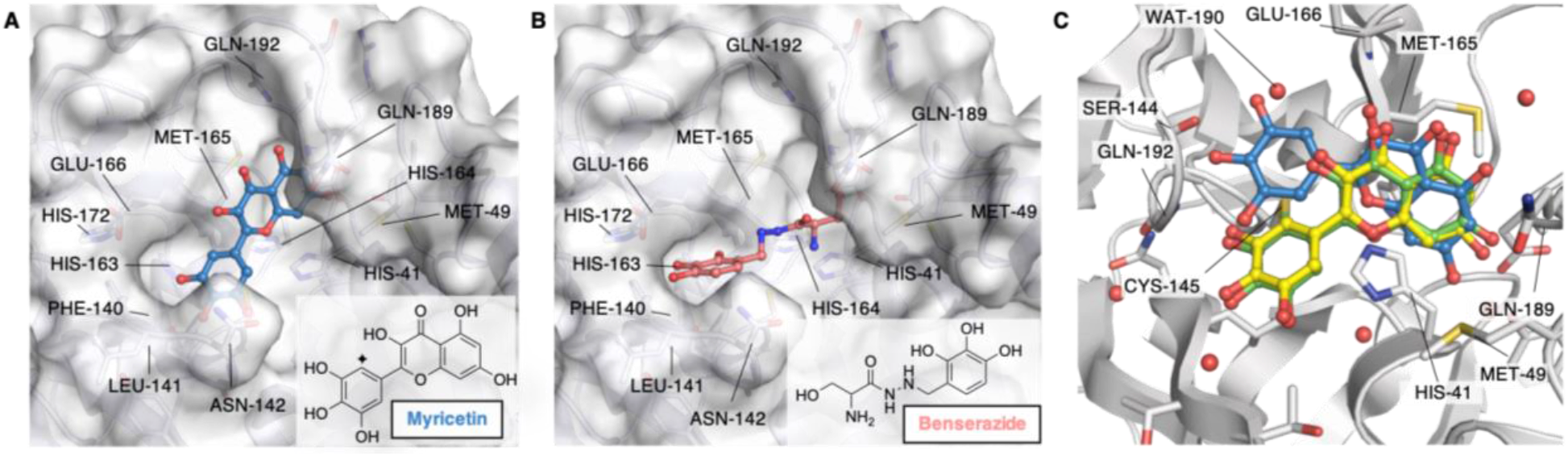
Binding poses of predicted high affinity ligands. Binding poses of Myricetin (EOS100914, **A**) and Benserazide (EOS100736, **B**), predicted to be high affinity binders using our active-pharmacophore model, and experimentally confirmed to be nM SARS-CoV-2 Mpro inhibitors. The protein structure is shown as white surface (PDB ID 6WTT, chain A), while Myricetin and Benserazide are shown in blue and coral ball-and-sticks, respectively. The poses shown here are the best scored ones according to the Dice coefficient. The insert panels show the molecular formulas of Myricetin and Benserazide. The diamond symbol in the scheme of panel **A** highlights the position of the nucleophilic attach by Cys145 on Myricetin. (**C**) Overlay of crystal structure (PDB ID 7B3E, green), docked (blue, RMSD 3.14 Å) and refined (yellow, RMSD 0.46 Å) binding poses of Myricetin. Binding pocket residues are shown in white ribbons and sticks with heteroatoms colored according to the atom type. The orientation of panel **C** was rotated with respect of those of panels **A** and **B** to show the covalent bond found in the X-ray crystal structure between Cys145 and Myricetin reactive carbon.

The docking pose of Myricetin, as coming out from our virtual screening procedure, shows an orientation which is comparable to the one observed in the newly solved X-ray structure with PDB ID 7B3E (resolution 1.77 Å, see Figure 6C). In this pose, the bicyclic ring in the two structures nicely overlap, while the 3,4,5-trihydroxyphenyl moiety is rotated in our predicted pose with respect to the crystallographic one. By refining the docking pose (see Section S9.1 and Figure 6C) both the bicyclic ring and the 3,4,5-trihydroxyphenyl moiety assumes an orientation nearly identical to the one found in the X-ray pose after covalent binding with Cys145, with an overall RMSD of 0.46 Å between the refined predicted pose and the crystallographic one. Interestingly, Baicalin features the same isoflavon scaffold as Myricetin, yet it binds the protein with a different orientation, as shown by X-ray studies (PDB ID 6M2N). Our procedure predicted such orientation, although the overall binding pose differed more significantly from the X-ray one than that with of Myriecitin (see Section S9.2, Figure S22).

Myricetin and Benserazide contain polyhydroxy-phenolic moieties, which are considered promiscuous due to their redox features but also to the presence of a high number of close Hbond acceptor/donor sites that allow them to satisfy several 3D-pharmacophores. Nonetheless, these compounds have, respectively, reached approved clinical usage (i.e. for Parkinson’s disease^82^ and alcohol use disorder^83^) and are in use in our diet like other polyhydroxyphenol-containing products.^84^ Also, Quercetin, structurally similar to Myricetin, was identified mild inhibitor of SARS-CoV-2 Mpro (K_I_ ~ 7 μM).^85^

## DISCUSSION AND CONCLUSIONS

SARS-CoV-2 Mpro is an important target for COVID-19 drug discovery because of its key role for viral replication and low similarity with human proteases.^6, 7^ Given its conserved nature with respect to the other Mpros across Coronaviruses and the presence of a huge number of crystallized structures (*apo* and *holo*), several drug repurposing and structure-based drug design campaigns have been conducted.^20–29^ Unfortunately, this has so far led to only 7 SARS-CoV-2 Mpro inhibitors in the nM range (IC50 ≤ 400 nM). This contrasts with SARS-CoV Mpro, for which 127 nM inhibitors are known in this range.^46, 47, 52, 54, 56, 60–62, 65^ The observed difficulties in identifying potent SARS-CoV-2 Mpro inhibitors was suggested to arise from the large plasticity of the binding site,^18^ along with other factors (also observed for SARS-CoV Mpro), including induced-fit conformational changes and formation of covalent bonds upon ligand binding^30^. Therefore, the available binding space can differ significantly from ligand to ligand.

Accounting for receptor binding site flexibility in molecular docking is a significant challenge. This can be partially overcome with a careful choice of the most appropriate receptor and reference ligand(s) or by performing ensemble docking approaches.^86, 87^ While it seems logical to employ multiple protein structures and ligands where available, very few published studies have systematically evaluated the impact of using additional information on proteins and ligands’ structure.^88^ These studies arrive at the conclusion that alternative structure-based design approach may be to define pharmacophores based on the binding site and use them to search large chemical databases.^89, 90^

Our paper exploits the particularly large amount of structural information available for Mpro (~200 X-ray structures in the *apo* and in the *holo* forms), along with a very long MD simulation from the D. E. Shaw group^37^ and structures generated here by enhanced sampling of the binding site dynamics. First, we have determined the potential druggability of each of the ~30,000 Mpro conformations generated from these sources, as calculated using TRAPP and SiteMap druggability tools.^38, 42, 43^ Next, we have used a sample library to understand how selected potential high-druggable protein conformations perform when probed with a diverse chemical space, here defined by 13,000 compounds (see t-SNE plot in Figure 7, and methods for details).

**Figure 7.**
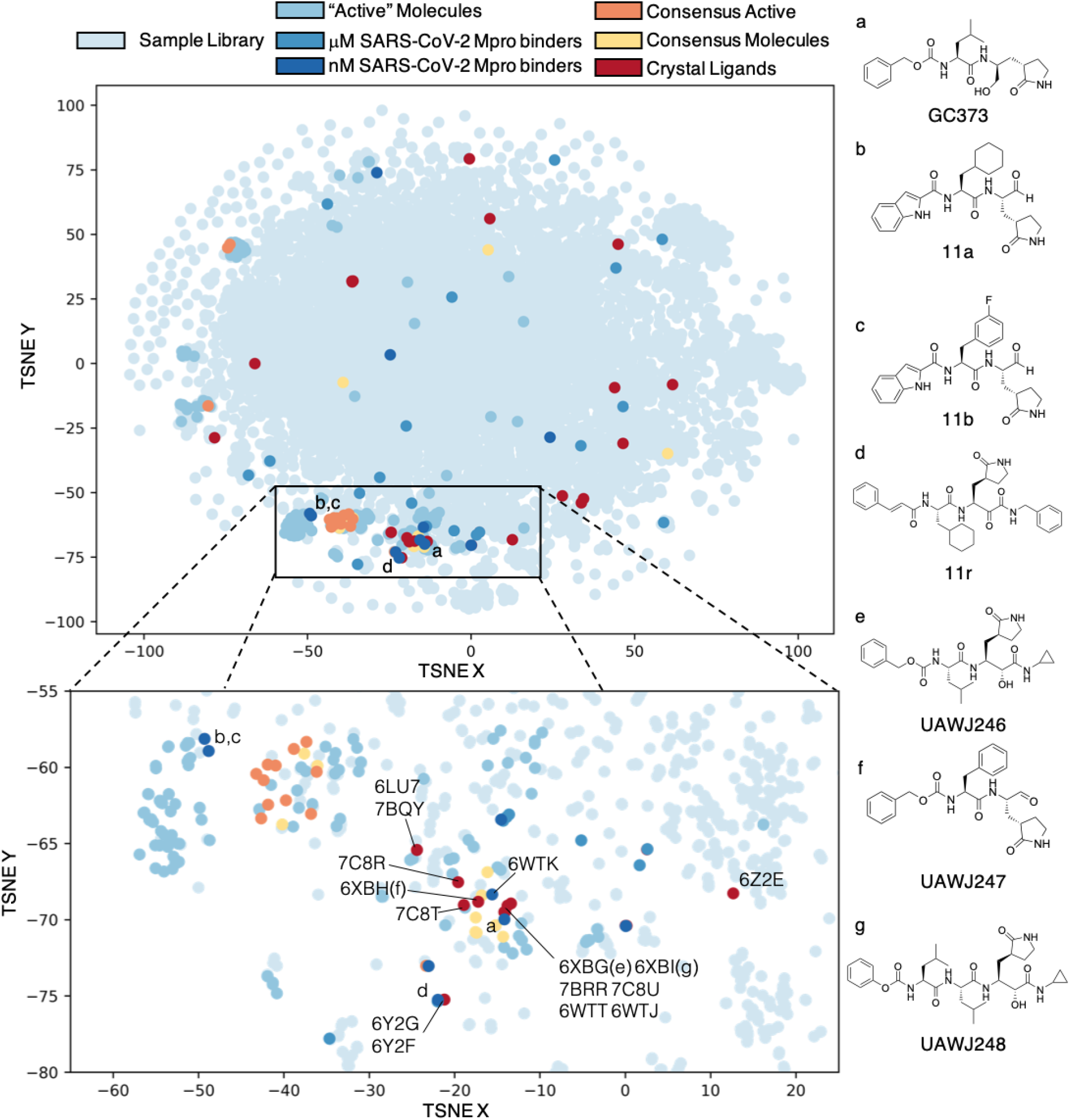
t-Distributed Stochastic Neighbor Embedding (t-SNE) plot of the sample chemical library screened (see Methods and SI Equation S4.2 for details). The sample library, the molecules denoted as “active” (due to their experimental binding affinity towards SARS-CoV), as well as the known SARS-CoV-2 Mpro binders are all plotted in different shades of blue. The 2D representations of a selection of nM affinity SARS-CoV-2 Mpro binders colored in the darkest blue shade (labeled a-d) are shown on the side. Ligands identified in the top 100 of the well-performing structures (“consensus”) are colored in yellow, while the subset of those that also have a high affinity towards SARS-CoV in experiments are plotted in orange. Lastly, the co-crystalized ligands are shown in red, with selected ligands shown in 2D representation on the side (labeled e-g). The inset with a magnified portion of the t-SNE plot is reported at the bottom of the figure. The “active” molecules appear to be more chemically diverse than the SARS CoV-2 Mpro co-crystallized ligands since they are spread over all the t-SNE plot, while the co-crystallized ligands mostly cluster in the bottom part of the plot. This region corresponds to peptides covalently bound to SARS-CoV-2 Mpro-C145 (PDB ids: 6LU7, 6LZE, 6M0K, 6WTJ, 6WTK, 6WTT, 6XA4, 6XBH, 6XBG, 6XBI, 6Y2F, 6Y2G, 6YZ6, 6Z2E, 7BQY, 7BRR, 7C8R, 7C8T, and 7C8U, see Table S1).

Our library included also “active” molecules, i.e. molecules that are known to bind with nM affinity (pIC50 > 6) to SARS-CoV Mpro. Therefore, virtual screenings against ~200 protein conformations were performed. We found that only a few of these highly-druggable (“well-performing”) conformations recognize a sufficiently high percentage of such “active” molecules: The ones in common among across the well-performing structure, “consensus active” molecules (16 in total) represent a small subgroup of the overall “active” molecules’ ensemble with specific features. In particular, the “consensus active” (16 molecules) are mostly clustered in two main areas of the t-SNE plot, both corresponding to peptidomimetic structures but differing from each other by the presence of a benzothiazole moiety and an additional peptide bond. The specific protein-ligand interaction fingerprint (PLIF) of the “consensus active” molecules strikingly resembles the one emerging from the SARS-CoV-2 Mpro co-crystalized ligands. The latter indeed cluster in the same region of the t-SNE plot. Notably, no consensus was found for the poorly-performing structures.

We then combined the crystallographic and the virtual screening PLIFs (i.e. the chemical space emerging from experimental structures and the chemical space selected upon virtual screening). Within the limitations of the procedure (discussed in the Limitation paragraph), we obtained an “active-pharmacophore” that we first used against a selection of SARS-CoV-2 Mpro binders (46 molecules): the latter are very diverse and they are spread overall the t-SNE plot (Figure 7). The active-pharmacophore could predict known nM-binders for SARS-CoV-2 Mpro (12 molecules out of the total of 46), which are also clustered in the peptides and peptidomimetics region of the t-SNE plot, and discriminate these from the μM ones. Moreover, it could also discriminate the transferable from the non-transferable binding features from SARS-CoV to SARS-CoV-2 Mpro.

The former include the interaction with the catalytic dyad residues along with: (i) His163, whose mutation to Ala inactivates SARS-CoV Mpro,^91^ (ii) Glu166, which plays a role in the dimerization (required for enzymatic activity) in SARS-CoV.^92^ In addition, its interactions with the N-finger of the other subunit assist the correct orientation of residues in the binding pocket for both proteins,^92, 93^ (iii) Gln189, which correlates evolutionally with residues from the Cys44-Pro52 loop in both proteins, which was shown to regulate ligand entrance to the binding site^18^ in both proteins, and (iv) Ser144, whose mutation to Ala hampers the catalytic activity in SARS-Cov MPro.^94^

The non-transferable binding features include the ability to place large hydrophobic/aromatic groups in the part of the cavity defined by Thr25 and Thr26 that is partially lost in SARS-CoV-2 Mpro compared to SARS-CoV Mpro. This appears to affect HBonds with Gly143 and Ser144. Accordingly, this cavity is empty or occupied by a smaller aromatic group like a benzyl ring in all the known nM-binders and the co-crystallized ligands of SARS-CoV-2 Mpro. In contrast, several of the known nM-binders of SARS-CoV Mpro have benzothiazole or analogous bulky aromatic groups in this position.

We finally used our active-pharmacophore against a public library of compounds. We predicted two ligands to be nM for SARS-CoV-2: Benserazide and Myricetin. Biochemical assay experiments confirmed them to be nM binders of SARS-CoV-2 (Figure S21). Our predicted pose of Myricetin is in very good agreement with the just solved X-structure of the complex (Figure 6C). This was not expected since Baicalin baring the same isoflavon scaffold of Myricitin binds in a reversed position (PDB ID 6M2N). Thus, the pharmacophore not only successfully predicts poses in a highly flexible binding site as that of SARS-CoV-2 Mpro, but it also discriminates between different orientations of quite similar scaffolds.

Our methodological approach also demonstrates that only a small fraction of the binding sites of the *apo* protein from crystal structures or simulations are similar to those of the well-performing *holo* structures. However, even the conformations with high structural similarity and high druggability scores, generated by molecular dynamics simulations, yielded low enrichment factors in virtual screening. Thus, druggability assessment methods fail to discriminate between small structural variations of the binding site that lead to successful performance in virtual screening. These small structural differences significantly impact ligand binding predictions as observed in our virtual screening campaigns. They could also be a source of disappointing results in other virtual screening campaigns carried out so far by research groups world-wide.^22, 28, 32, 33, 95^ These observations indicate that there is space to improve the discriminatory ability of druggability scores by training on a wider range of structures generated by simulation as well as crystallography. Moreover, they highlight the need to develop simulation methods to generate *holo*-like protein structures for virtual screening.

This work was carried out within the framework of EXSCALATE4CORONAVIRUS (E4C)^79^ project. E4C aims to exploit the most powerful computing resources currently based in Europe to empower smart in silico drug design applied to the 2019-2020 SARS-CoV-2 coronavirus pandemic, while increasing the accuracy and predictability of Computer-Aided Drug Design (CADD). We here used 400,000 core-hours on the JURECA supercomputer in the Jülich Supercomputing Centre. Advanced CADD in combination with high throughput biochemical and phenotypic screening are allowing the rapid evaluation of simulation results and the reduction of time for the discovery of new drugs.^96^ The work presented here indeed shows how a computational procedure combined with experimental validation can correctly predict structure and affinity trends of effective hit molecules for a given target. This kind of combined approaches may be a key strategy especially against pandemic viruses and other pathogens, where the immediate identification of effective treatments is of paramount importance.

## EXPERIMENTAL PROCEDURES

### Expression and enzymatic activity

SARS-CoV-2 Mpro (ORF1ab polyprotein residues 3264-3569, GenBank code: MN908947.3) has been produced, purified and used as described in Zhang *et al.*^17^ The detection of enzymatic activity of the SARS-COV-2 3CL-Pro was performed under conditions similar to those reported by Zhang *et al.*^17^ Enzymatic activity was measured by a Förster resonance energy transfer (FRET), using the dual-labelled substrate, DABCYL-KTSAVLQ↓SGFRKM-EDANS (Bachem #4045664) containing a protease specific cleavage site after the Gln. In the intact peptide, EDANS fluorescence is quenched by the DABCYL group. Following enzymatic cleavage, generation of the fluorescent product was monitored (Ex 340 nm, Em 460 nm), (EnVision, Perkin Elmer). The assay buffer contained 20 mM Tris (pH 7.3), 100 mM NaCl and 1 mM EDTA. The assay was established in an automated screening format (384 well black microplates, Corning, #3820)

### Primary screen and dose response

In the primary screen, test compounds (stock at 10 mM in 100 % DMSO), positive (Zinc-Pyrithione [medchemexpress, Cat. No.: HY-B0572] 10 mM in 100 % DMSO) and negative (100 % DMSO) controls, were transferred to 384-well assay microplates by acoustic dispensing (Echo, Labcyte). Plate locations were: test compounds at 20μM final (columns 1 to 22); positive control Zinc-Pyrithione at 10 μM final (column 23); and negative control 0.2 % v/v (column 24). 5 μl of SARS-CoV-2 Mpro stock (120 nM) in assay buffer were added to each plate well and incubated with the compounds for 60 min at 37 °C. After addition of 5 μl substrate (30 μM in assay buffer), the final concentrations were: 15 μM substrate, 60 nM SARS-CoV-2 Mpro, 20 μM compound, and 0.2% DMSO, in a total volume of 10 μL/well. The fluorescence signal was then measured at 15 min and inhibition (%) calculated relative to controls. To flag possible optical interference effects fluorescence was also measured 60 min after substrate addition, when the enzymatic reaction was complete. Results were normalized to the 100 % and 0 % inhibition controls.

### Hit follow up in confirmation, profiling and counter assays

For Hit Confirmation (HC), the compounds were re-tested in the same primary assay format in triplicate at 20 μM compound concentration, final. Confirmed compounds were then profiled in triplicate in 11 point concentration responses, starting from 20 μM top concentration with 1:2 dilution steps. To flag optical and non-specific interference effects, a counter screen was performed during hit profiling with the assay protocol adjusted so that compound addition occurred at 60 min post substrate addition, when the enzymatic reaction was complete.

### Markov State Modeling

Kinetically distinct states were identified from the 100 *μ*s simulation^37^ from D. E. Shaw Research through Markov state modeling. Two alternative Markov state models were created with two different sets of input features 1) with Cartesian coordinates of all alpha carbons in the dimer and 2) 10,668 distances in the range 0.5 to 1.3 nm spread across the whole dimer. Further dimensionality reduction was done with time-lagged independent component analysis (tICA)^97, 98^ in five dimensions and with a lag time of 10 ns. The obtained 5-dimensional space was further discretized into 50 distinct states through k-means clustering, which were used to construct Markov state models with lag time 10 ns. Perron cluster-cluster analysis, PCCA++, was finally used to obtain 1) three and 2) four kinetically distinct macrostates. Markov state modeling was done with the software PyEMMA^99^ and visualizations with VMD.^100^

### Druggability analysis of Sars-CoV2 main protease structure

The SiteMap tool,^101^ together with the TRAPP (TRAnsient Pockets in Proteins) approach^42^ were used for the characterization of the proteins’ binding sites. The SiteScore is based on a weighted sum of several properties accounting for the degree of pocket enclosure (a surrogate for pocket curvature), pocket size, and the balance between hydrophobic and hydrophilic character in the binding site. The Dscore uses almost the same properties as SiteScore but different coefficients are used and hydrophilicity is not considered. TRAPP provides tools for (i) the exploration of binding pocket flexibility and (ii) the estimation of druggability variation in an ensemble of protein structures.^42^ Specifically, two methods were applied: Langevin Rotationally Induced Perturbation (LRIP)^39^ and tConcoord^102^ to generate conformational ensembles that explore the flexibility of the Mpro binding pocket. Subsequently, the binding pocket’s druggability in each of the generated protein conformations was computed (using LR and CNN methods,^41^ see below for more details). Additionally, we computed the druggability for 10,000 snapshots taken at 10 ns intervals from the 100 μs standard MD trajectory generated by starting from the crystal structure with PDB ID 6Y84 by D. E. Shaw Research.^37^

### Generation of binding pocket conformational ensembles using LRIP and tConcoord

We based the conformational ensemble generation on the high-resolution X-ray crystal structures with PDB IDs 6LU7 (2.16 Å resolution^103^) and 6Y2G (2.20 Å resolution^17^). All 306 residues of SARS-CoV-2 Mpro are resolved in PDB ID 6LU7; for PDB ID 6Y2G, the unresolved C-terminal residues (see Table S1) were modeled using PyMol.^104^ All TRAPP analysis was conducted on homodimeric structures, generated with the symmetry wizard of PyMol.^104^ All heteroatom records were removed from the protein structures and hydrogen atoms were added at a pH of 7.4 using the CHARMM force field^105^ and pdb2pqr.^106^ The structure of the covalent ligand (N3), needed to define the binding pocket in TRAPP, was obtained from the PDB ID 6LU7 and, in the case of 6Y2G, positioned in the binding pocket by alignment of 6LU7 to 6Y2G using PyMol^104^. For both crystal structures, 6LU7 and 6Y2G, 200 energy minimized conformations were generated with tConcoord^102^ using [TRAPP4](http://trapp.h-its.org/). Additionally, ensembles were generated with LRIP^39^ at 300 K, generating 100 conformations for each pocket lining residue. For 6LU7, 100 perturbations were made, each followed by 100 MD steps for relaxation. For 6Y2G, 300 perturbations were made, each followed by 300 MD steps for relaxation. The parameters for 6Y2G were chosen to increase conformational sampling and improve sampling statistics.

### Druggability calculation

The active site pocket of SARS-CoV-2 Mpro was defined with TRAPP4 by assigning a distance of 3.5 Å around all atoms of the ligand N3 from PDB ID 6LU7. This distance was used to detect residues that potentially may contribute to the binding site and to define dimensions of a 3D grid that was then used to compute the binding pocket shape. Then the binding pocket for each structure was mapped on the 3D grid. The druggability score of this pocket was computed using linear regression (LR) or a convolutional neural network (CNN) and scaled between 0 and 1.^41^ Scores were calculated for all conformations generated for 6LU7 and 6Y2G from LRIP and tConcoord, as well as for frames collected every 10 ns from the conventional MD simulations.^37^ For each structure, a set of the binding site residues that line the binding pocket was detected using the procedure implemented in the TRAPP package. Specifically, each residue was characterized by the number of atoms that contact with the binding pocket.

Structure Selection: First we selected all structures (generated by tConcoord, LRIP, or extracted from MD trajectories) with the top 10% of the LR and CNN druggability score. These structures were then clustered by the binding site similarity into 8 clusters using k-means procedure (see Figure S8 and also Section S4.3 for more details).

### Library Preparation

Virtual screening studies were performed on a repurposing library including all the commercialized and under development drugs retrieved in the Clarivate Analytics Integrity database, merged with the internal chemical library from Dompè pharma company of already proven safe in man compounds and the Fraunhofer Institute BROAD Repurposing Library, removing duplicate structures. Known inhibitors of SARS-CoV Mpro, retrieved from several sources, literature, the Clarivate Analytics Integrity database, the GOSTAR database and the data repository shared by the Global Health Drug Discovery Institute, were added. A final unique list of about 13,500 drugs were obtained. All compounds^46, 47, 52, 54, 56, 60–62, 65^ were converted from 2D to 3D and prepared with Schrödinger’s LigPrep tool.^107^ This process generates multiple sets of coordinates for different stereoisomers, tautomers, ring conformations (1 stable ring conformer by default) and protonation states. The Schrödinger Epik software^108, 109^ was used to assign tautomers and protonation states that would be more populated at the selected pH range (pH = 7 ± 1). Ambiguous chiral centers were enumerated, allowing a maximum of 32 isomers to be produced for each input compound. OPLS3 parameters were generated for each ligand. Multiple conformations for each compound were generated and a 1,000 step torsional sampling was performed. The conformers were retained if the minimized energy of the conformer is within 50 kJ/mol of the global minimum.^110^ After preparation, this resulted in 23,000 coordinate files for different conformers, tautomers and protomers of the compounds in the library.

### Glide Docking

The protein was preprocessed with the Protein Preparation Wizard from the Schrödinger Suite version 2019-4 with the default parameters.^108, 109, 111, 112^ The protonation states of each side chain were generated using Epik for pH = 7 ± 2.^108, 109^ All water molecules were removed. Energy minimization was performed using the OPLS3 force field.^110, 113^ Glide Version 85012^69, 70^ was used for all docking calculations. Docking of the compounds, prepared as described above, was performed to both active sites of the homodimer. The grids for the docking were prepared using the default parameters, with the internal grid box centered on the centroid of the co-crystallized ligand, when available. The external grid box was defined by checking the option “Dock ligands similar in size” (~32 x ~32 x ~32 Å). The internal grid box guides the docking algorithm to the region of interest, while the external grid box allows greater flexibility in ligand size and orientation. The Glu166 (backbone nitrogen bound hydrogen atom as well as the backbone oxygen atom), His163 (sidechain NE2 bound hydrogen atom) and His164 (sidechain oxygen atom) were defined as possible Hbond constraints. A standard precision (SP) Glide docking was carried out, generating 20 poses per docked molecule; During the docking procedure, poses were rejected, if they were not able to fulfill at least 2 of the Hbond constraints defined above. Glide score version 5.0^114^ was used to rank the binding poses. This is an empirical scoring function designed to reproduce trends in the binding affinity.

### FRED Docking

The SARS-CoV-2 Mpro receptors were generated by OpenEye Spruce4Docking,^68, 115, 116^ using as input the structures pre-processed by the Protein Preparation Wizard and Epik in the Schrödinger Suite. The protonation state of the receptor was not altered. The OpenEye Docking suite performs rigid docking of pre-generated conformers. Here, these conformers were generated from the prepared library (see above) using OpenEye OMEGA.^117^ Conformers with internal clashes and duplicates were discarded by the software and the remaining ones were clustered on the basis of the root mean square deviation (RMSD). For virtual screening, a maximum of 200 conformers per compound, clustered with a RMSD of 0.5 Å, was used. If the number of conformers generated exceeded the specified maximum, only the ones with the lowest energies were retained. Rigid-body docking was performed using OpenEye FRED,^68, 115, 116^ which is included in the OEDocking 3.4.0.2 suite.^68, 115, 116^ Each conformer was docked by FRED in the negative image of the active site of the target protein, which consists of a shape potential field in the binding site volume. The highest values in this field represent points where molecules can have a high number of contacts, without clashing into the protein structure. In its exhaustive search, FRED translates and rotates the structure of each conformer within the negative image of the active site, scoring each pose. The first step has a default translational and rotational resolution of 1.0 and 1.5 Å, respectively. The 100 best scoring poses were then optimized with translational and rotational single steps of 0.5 and 0.75 Å, respectively, exploring all the 729 (six degrees of freedom with three positions = 36) nearby poses. The best scoring pose was retained and assigned to the compound. The binding poses were evaluated by using the Chemgauss4 scoring function implemented in OpenEye.^68, 115, 116^

### Molecular Fingerprint generation

Chemical circular fingerprints used for the t-SNE plots in this work were generated using the RDKIT package in an in-house python script.^118^ Circular and dictionary type fingerprints (ECFP4/6, Molprint2D and MACCs) used for database diversity calculations were generated using Schrödingers Canvas program.^118–120^

PLIF generation and representation. For the generation of the Protein Ligand Interaction Fingerprints (PLIF) the Open Drug Discovery Toolkit (ODDT) package was implemented in an in-house python script.^121^ Each PLIF represents a group of docked or co-crystallized molecules saved as a series of binary fingerprints, each one representing the interaction of the molecule itself with the receptor residues. For each residue of the receptor, 8 bits are used to describe the presence or absence of different types of interaction (hydrophobic, aromatic face-to-face, aromatic edge-to-edge, Hbond acceptors and donors, and ionic interactions) with the ligand. The active-pharmacophore was derived by the average of the fingerprint of the “active” molecules in the consensus, docked on the well-performing structures, with the fingerprint of the co-crystallized ligands in the X-ray structures considered in this work. The resulting vector components were then rounded according to a cutoff in order to form a bit vector representation. The most suitable cutoff value was derived by iterating cutoff values between 0.1 and 0.9 and calculating the Dice similarity index between the resulting bit vector and the PLIF generated by the docking poses of the known SARS-CoV-2 Mpro binders. The cutoff, which would result in the highest dice similarity when compared to the nanomolar affinity ligands was chosen. The molecules used for generation of the crystal structure consensus fingerprint slightly overlap (7 out of 36 crystal ligand conformations) with the sub-μM affinity ligands that are scored with the dice coefficient. When removing these from the query, the pharmacophore is still able to locate the higher affinity ligands in the top positions. The same Dice was screened against a selected number of EU-OPENSCREEN ERIC Bioactive Compound Library. The PLIF representation in the form of stacked histograms used in this work was generated using python matplotlib library (v 3.3.1).^122^ In the histograms, each interaction is represented by a bar, whose height is proportional to the number of times the interaction was observed among the group of molecules, divided by the number of molecules. Bars representing different types of interactions with the same residue are stacked onto each other, to visualize more easily the most important residues.

T-SNE plot. T-Distributed Stochastic Neighbor Embedding (t-SNE) plot. Embedding is based on the 2048-bit Morgan^118^ fingerprint with a radius of 3. Scikit-learn t-SNE implementation with a custom Tanimoto distance metric was used (see supporting information) in an in-house python script. This algorithm performs a nonlinear dimensionality reduction technique that models each high-dimensional object, i.e. a molecule in this case, by a two-dimensional point, in such a way that similar objects (i.e. similar molecules) are modeled by nearby points and dissimilar objects are modeled by distant points.

### Limitations

As with any modelling study, our models also have limitations. Protein mobility is most probably increased by the presence of moving and displaceable water molecule ^123^. In the molecular screenings, individual water molecules, which may be critical in the binding process, were not accounted for. Solvation effects can account for up to 100-fold difference in binding affinity (corresponding to ~3 kcal/mol in binding free energy^124^). Also, we in part rely on scoring functions to rank and select the best binding poses. Current docking/scoring methods^124, 125^ were suggested to provide reasonable predictions of ligand binding modes, but their performance is often disappointing in predicting ligand binding affinities. Additionally, those methods are often system-dependent, making it very hard to decide which scoring function is suitable for the chosen target protein. To partially overcome this issue, we carefully set-up Glide and Fred docking procedure by reproducing a set of covalent and non-covalent SARS-CoV-2 Mpro-ligand crystal poses.

Moreover, we compare assay-dependent IC50 data coming from different laboratories. In addition, several of the known Mpros inhibitors are covalently bound to the protein. Irreversible (covalent) enzyme inhibitors cannot be unambiguously ranked for biochemical potency by using IC50 values, because the same IC50 value could originate either from relatively low initial binding affinity accompanied by high chemical reactivity, or the other way around^126^. In other words, the important quantity to be considered would be the rate of covalent modification, (k_inact_/K_I_), that describe the efficiency of covalent bond formation resulting from the potency (K_I_) of the first reversible binding event and the maximum potential rate (k_inact_) of inactivation.^127^ This information is unfortunately not available for most of the ligands here considered.

## Resource availability

Further information and requests for resources should be directed to and will be fulfilled by the lead contact, Giulia Rossetti.

## Materials availability

All materials generated in this study are available from the lead contact without restriction.

## Data and code availability

The datasets generated during this study are available at DOI: 10.5281/zenodo.4299967

## SUPPLEMENTAL INFORMATION

Supplemental experimental procedures, Figures S1–S22, and Table S1-S6. Table S1-3 are available at DOI: 10.5281/zenodo.4299967

## ACKNOWLEDGMENTS

Additional computational resources were provided by the Swedish National Infrastructure for Computing (SNIC) and the Knut and Alice Wallenberg Foundation. D.B.K and R.C.W. acknowledge the support of the Klaus Tschira Foundation. G.R., P.C., D.B.K and R.C.W. acknowledge the Human Brain Project founded by the European Union’s Horizon 2020 Framework Programme for Research and Innovation under the Specific Grant Agreement No. 945539 (Human Brain Project SGA3). G.R. A.Z., P.S. and P.C. acknowledge the E4C consortium. We would also like to thank Dr. Katja Herzog (EU-OPENSCREEN ERIC) for indicating access to the EU-OPENSCREEN ERIC Bioactive Compound Library data^80^.

## AUTHOR CONTRIBUTIONS

J.G. and S.A. performed all the docking experiments, the PLIF and pharmacophore calculations, as well as editing most of the figures and tables of the manuscript. A.H. and D.B.K. performed the TRAPP analyses. B.P.J. collected and analyzed the X-Ray structures. F.M. and G.R. performed the SiteMap analyses. C.M., C.T. and A.B. took care of the library collection. C.B. and E.L. performed the MSM analyses. M.K., P.G. and A.Z. performed the experiments. E.C. and P.S. solved the crystal structure. F.S., A.Z., P.C., R.C.W., F.M., D.B.K. and G.R. wrote the manuscript and contributed to the design of the research and the analysis of the data.

## DECLARATION OF INTERESTS

The authors declare no competing interests.

## Notes

https://doi.org/10.5281/zenodo.4299967

### Competing Interest Statement

The authors have declared no competing interest.

### Summary of Updates

Correction in author name

