## Supporting Information for "A blueprint for high affinity SARS-CoV-2 Mpro inhibitors from activity-based compound library screening guided by analysis of protein dynamics"

### SECTION S1. Mpro conservation.

We report here the sequence and structural comparison between SARS-CoV and SARS-CoV-2 Mpro enzymes (Figure S1).

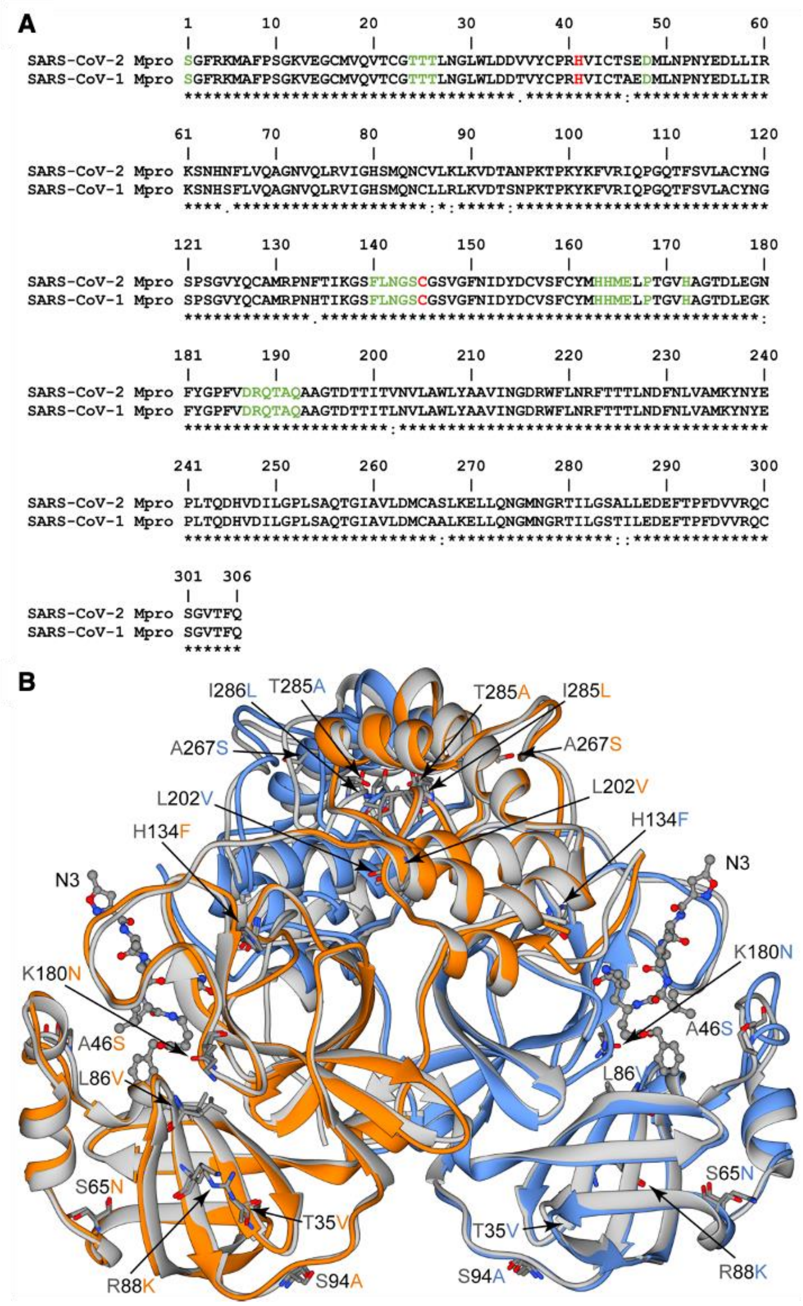


**Figure S1. Mpro conservation.** (**A**) Sequence alignment of SARS-CoV and SARS-CoV-2 Mpro. The catalytic dyad is highlighted in red, and active site residues (detected considering the residues within 4 Å of the ligand N3 in the PDB ID 6LU7) are highlighted in green. (**B**) SARS-CoV Mpro dimer in *apo* form (PDB ID: 2DUC, in grey) superimposed with the corresponding protein from SARS-CoV-2, bound to the N3 covalent inhibitor (PDB ID: 6LU7, in orange and light blue). The N3 ligand is shown in ball-and-stick representation, while the residues that are not conserved are shown in stick representation colored according to atom-type.

**SECTION S2. Analysis of position-featured MSM.**

We report here the Markov State Model (MSM) analysis performed on the trajectory made available by D. E. Shaw Research.[1]

In detail, we used the MD trajectory to create a Markov state model (MSM), whose eigenvectors represent various dynamical processes organized according to the corresponding timescales (see Section S2, Figure S2 and S3). From these eigenvectors, two sets of kinetically distinct macrostates, separated by the highest energy barriers sampled in the simulation, could be obtained (Figure S2A). This was done by using 1) Cartesian coordinates of the alpha carbons as features, and 2) a large set of short distance features, followed by time-lagged independent component analysis (tICA) for dimensionality reduction and k-means clustering (see Methods for details and Section S2). By randomly sampling conformations from these two sets of macrostates, we obtained 30 and 40 representative conformations for models 1) - three-state model - and 2) - four-state-model, respectively (Table 1).

By looking at the eigenvector components of the tICs (Figure S2A,C), the first tIC mainly represents a hinging motion between the two subunits, while the second tIC is dominated by a few residues in one of subunit B. In the B subunit timescales were longer than in A, therefore the analysis was focused on the slower moving subunit B.

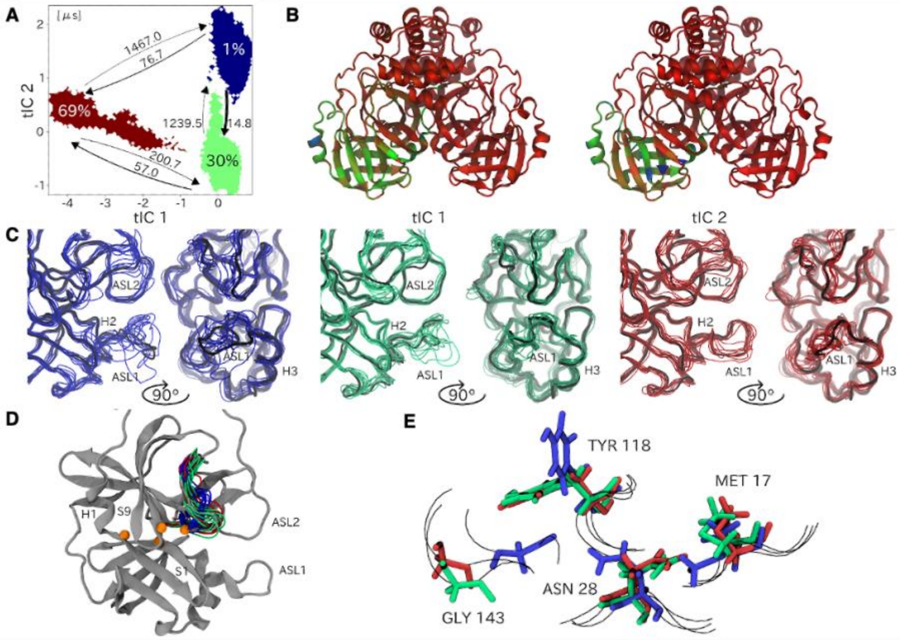


**Figure S2. MSM analyses.** (**A**) Three macrostates obtained from the MSM with position features, plotted onto the first two tICs. Percentages represent macrostate populations, arrows depict transition rates with numbers representing mean first passage times in microseconds. (**B**) Eigenvector components of the first two tICA coordinates. Blue colors represent a high contribution to the coordinate while red represents a low contribution. (**C**) The binding site of ten samples from each macrostate in panel (**A**). The black structure shows a representative conformation. Especially note differences in the conformation of ASL1 and H2. (**D**) Differences in the conformation of the loop adjacent to ASL2 between the blue state and the red and green states. Through analyzing residues with higher contributions to the second tIC, differences in four residues (orange) could be found. (**E**) Four residues that adopt different side-chain conformations in the blue macrostate. G143 sits on the displaced loop.

The macrostates were able to distinguish some differences around the binding site (Figure S2C), mainly in the loop ASL1 and helix H2. The least-populated blue state showed a very flexible ASL1 that fluctuated between open conformations and more closed ones, mainly seen in the green macrostate. H2 also moved between folded and unfolded states. In the green state, with a population of 30%, the ASL1 loop folded into a helix most of the time but occasionally remained in a flexible open conformation. H2 appears to generally be unfolded, as this is necessary in order to fold the ASL1 loop. The most populated red state corresponds to a rather stable closed conformation of the ASL1 loop, while H2 is stably folded. Another difference can be found in the second loop which is consistently turned away from the binding site in the blue macrostate in comparison to the green and red macrostates (Figure S2D). By analyzing some of the residues that had a larger contribution to the second tICA coordinate, we found differences in interaction patterns between residues Met17 (right between H1 and S1) and Asn28 (on S1) which both flip 180°, and residues Gly143 (on the displaced loop) and Tyr118 (on S9) that flip 90° (Figure S2E). In addition, other surrounding residues that did not exhibit a radical change in conformation between macrostates are most likely still involved in stabilizing these interactions.

**SECTION S3. Binding site analyses.**
The section is divided in three subsections: S3.1 where details on the analyzed Mpro binding sites' features are provided. S3.2 where the crystal packing of the deposited crystal structures is analyzed. S3.3 where the druggability details of the analyzed binding sites are provided. S3.4 where selection criteria for the MD10000 and tC/LRIP ensembles are provided.

**SECTION S3.1. Details on the features of the Mpro binding sites**

**Table S1. Available structures of SARS-CoV-2 Mpro selected for binding site analyses.** Available at DOI 10.5281/zenodo.4299967.

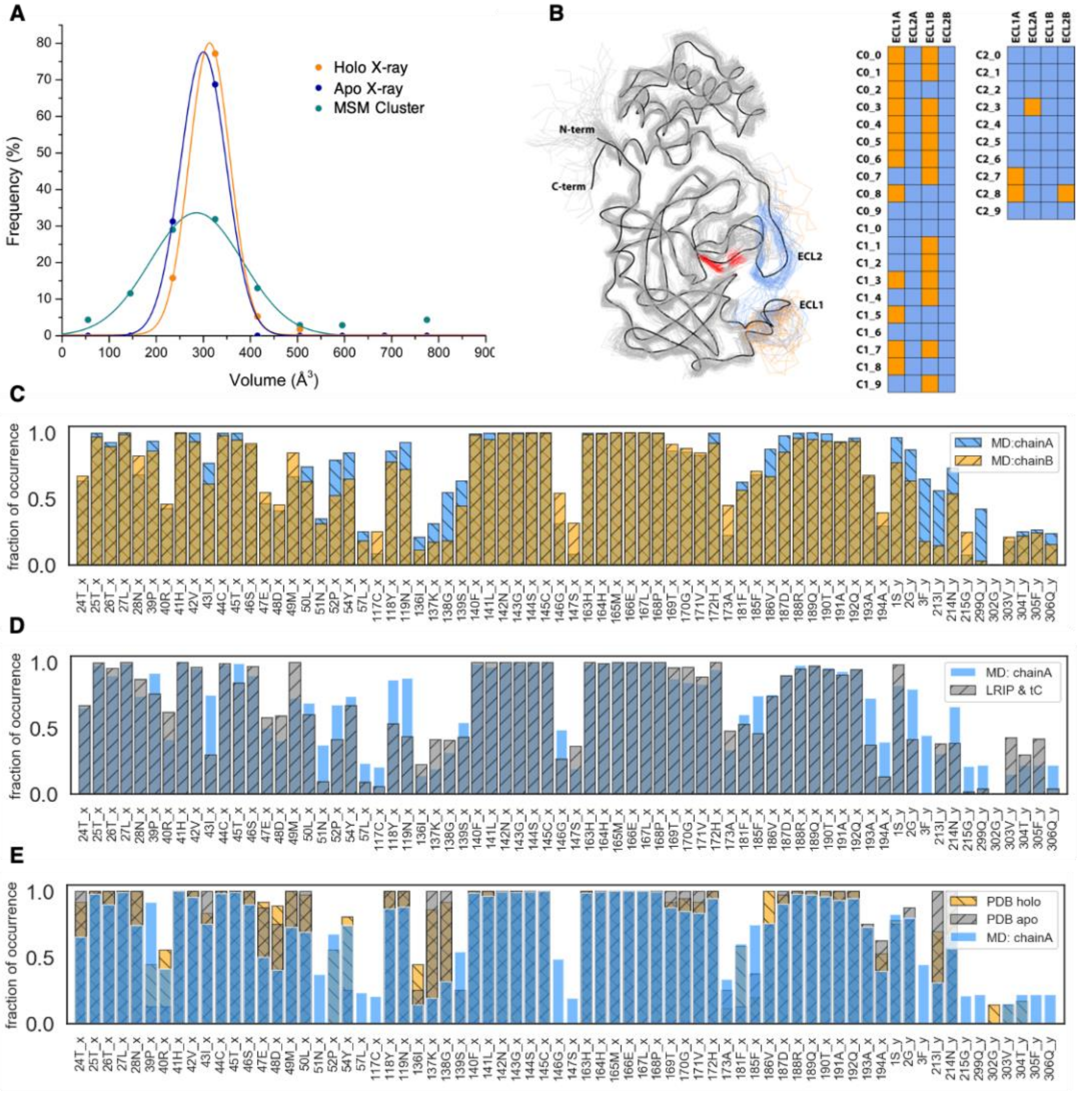


**Figure S3. Binding Site Analyses.** (**A**) Distribution of binding site volume, as calculated with the SiteMap tool, for the deposited X-ray crystal structures and the MSM cluster representatives. (**B**) Analysis of the conformation assumed by ASL1 and ASL2 (in both monomers) in the three-state Markov state model as derived from the µs-long trajectory published by D.E. Shaw Research team. Orange and blue squares refer to open and close conformations, respectively (**C**) Residues found in the binding sites during the µs-long trajectory published by the D.E. Shaw Research team. (**D**) Residues found in the binding sites for the LRIP/tC ensemble compared with those in the MD trajectory shown for chain A; and (**E**) residues found in the binding sites for the set of *apo*- and *holo*- crystal structures in the PDB and compared with those in the MD trajectory shown for chain A. x and y after the residue's number indicate two different chains ("x" denotes the chain whose binding site was analyzed).

**Loop Flexibility.** The structural fluctuations occur on the time-scale from nanoseconds (e.g. residues 47-54 in the ASL1 and 303-306 in the C-terminal tail of the chains A and B, respectively) to microseconds (e.g. residues 173-185 and 190-195 in the ASL2, chain A), confirming the higher flexibility in ASL1 vs. ASL2, as also emerged from the analysis of the MSM cluster representatives (Figure 2B and Figure S2-S3). After about 95 µs of MD simulation, we observed the folding of ASL2 to form a short helix in chain B only (see Figure 2C for chain B, and also Figure S4), which is also accompanied by a displacement of the loop consisting of residues 132-137 (tough the latter do not directly contribute to the active site).

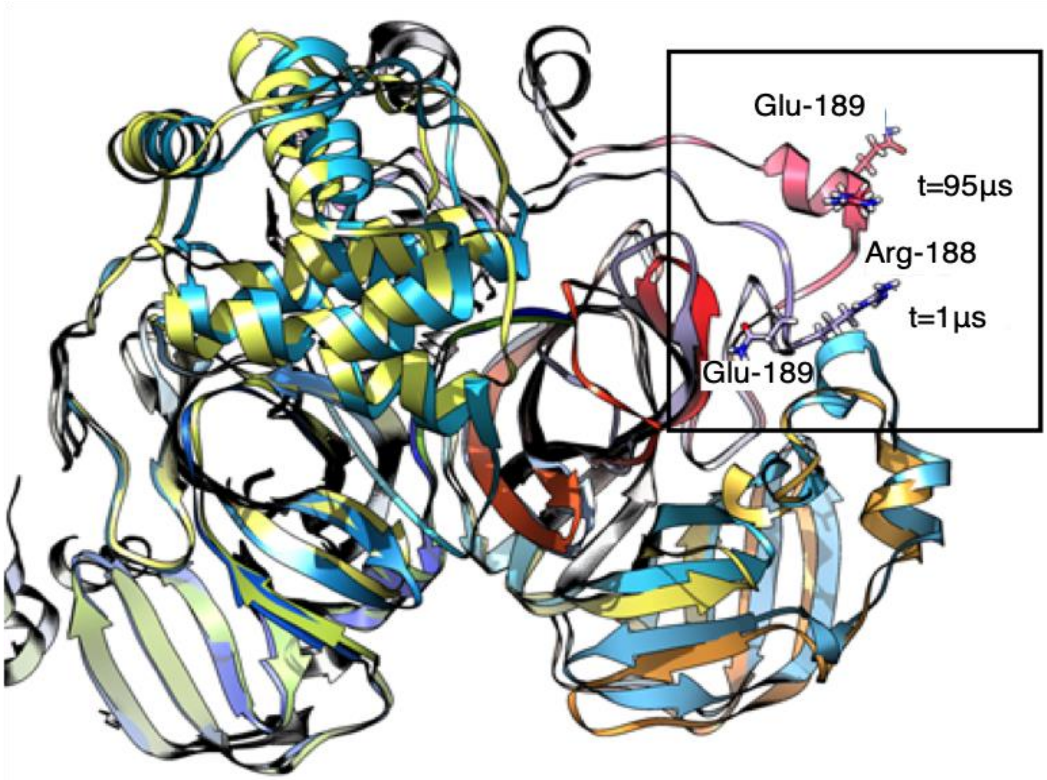


**Figure S4. Loop Flexibility.** Illustration of structural variations of the binding site at chain B observed after 95 µs of MD simulation: the loop A173-A194 transforms into a short helix (the corresponding region is shown by a rectangle); structures after 1 and 95 µs of MD simulation are shown in blue-green-violet and red-orange-yellow pallets, respectively.

Accordingly, in MSM analysis, this motion dominated the second tlC. This indicates that even 100 µs MD sampling is not sufficient to reveal the entire conformational space of the SARS-CoV-2 Mpro active sites.

**Table S2A. SiteScore analysis of all the deposited X-ray crystal structures for the Mpro.** Available at DOI 10.5281/zenodo.4299967.

**Table S2B. SiteScore analysis of the MSM ensemble (4-macrostates).** Available at DOI 10.5281/zenodo.4299967.

The X-ray crystal structures, M10000 and LRIP/tC TRAPP-pocket simulations are available here: DOI 10.5281/zenodo.4299967. Please note that there is a large variation of the druggability index in the LR model, while the CNN druggability index correlates better with that obtained from DScore and SiteScore. From nine structures with SiteScore > 1 (PDB IDs 5REJ, 5RFS, 5RFW, 6LU7, 6M2N, 6W63, 6WNP, 7BQY, and 7BUY), six also have CNN scores above 0.95.

**SECTION S3.2. About crystal packing.**

We checked the crystal packing of Mpro deposited structures. Mpro was crystallized in eight different space groups P $2_1$ $2_1$ $2_1$, P 1 $2_1$ 1, P 1, I 1 2 1, C 1 2 1, P $6_1$ 2 2, P $3_2$ 2 1 and P $2_1$ $2_1$ 2. For one representative structure of each space group, we calculated the minimum distance between two images, as well as the minimum distance between the binding site residues and the nearest image (see Table S3). For some space groups the packing is very tight, possibly stabilizing compact conformations of the binding site. While for others, the binding site is sufficiently far from the surrounding images to not be affected.

**Table S3. Minimum distance between non-hydrogen atoms in two Mpro images in selected structures for each space group in which Mpro was crystallized.** The minimum distance between the binding site residues and the closest Mpro image is also indicated.

| PDB ID | Space Group | Minimum Distance [Å] | Minimum Distance in the Bind Site [Å] |
|---|---|---|---|
| 6XHU (*apo*) | P 1 $2_1$ 1 | 2.63 | 7.73 |
| 7BQY (*holo*) | C 1 2 1 | 9.24 | 9.24 |
| 6W63 (*holo*) | P $2_1$ $2_1$ 2 | 3.11 | 7.09 |
| 6WQF (*apo*) | I 1 2 1 | 2.41 | 2.41 |
| 6WTT (*holo*) | P $3_2$ 2 1 | 4.07 | 4.07 |
| 6XBI (*holo*) | P 1 | 3.69 | 4.07 |
| 7C8R (*holo*) | P $6_1$ 2 2 | 3.51 | 5.09 |
| 6Y2G (*holo*) | P $2_1$ $2_1$ $2_1$ | 3.57 | 4.63 |

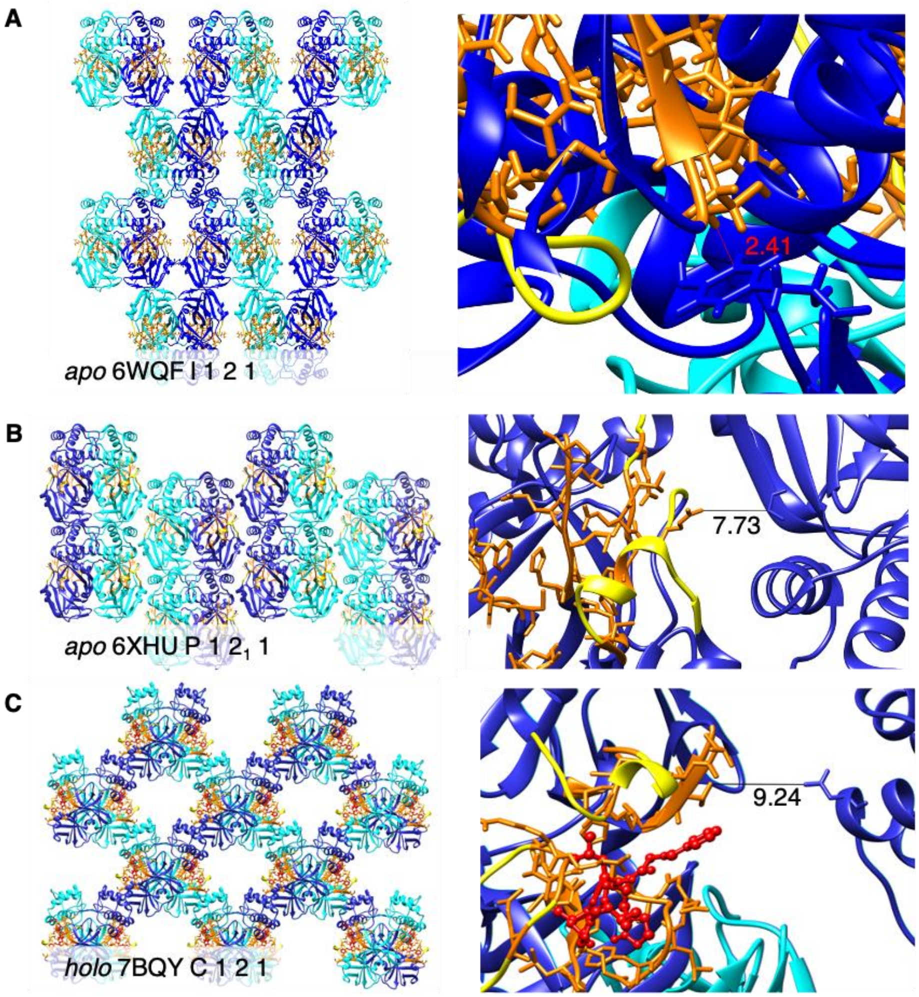


**Figure S5. Crystal packing of SARS-CoV-2 Mpro structures.** The analysis was done using the Unit Cell tool available in UCSF Chimera 1.14. The minimum distances between the binding site of one symmetry image and the atoms in the closest image for (**A**) 6WQF (*apo*), (**B**) 6XHU (*apo*) and (**C**) 7BQY (*holo*) are shown. Chain A, chain B and the floppy regions of the protein are shown as cyan, blue and yellow ribbons respectively. The 23 binding site residues are shown as orange sticks and the ligand is shown as red balls-and-sticks. All distances are in Å.

### SECTION S3.3. Druggability.

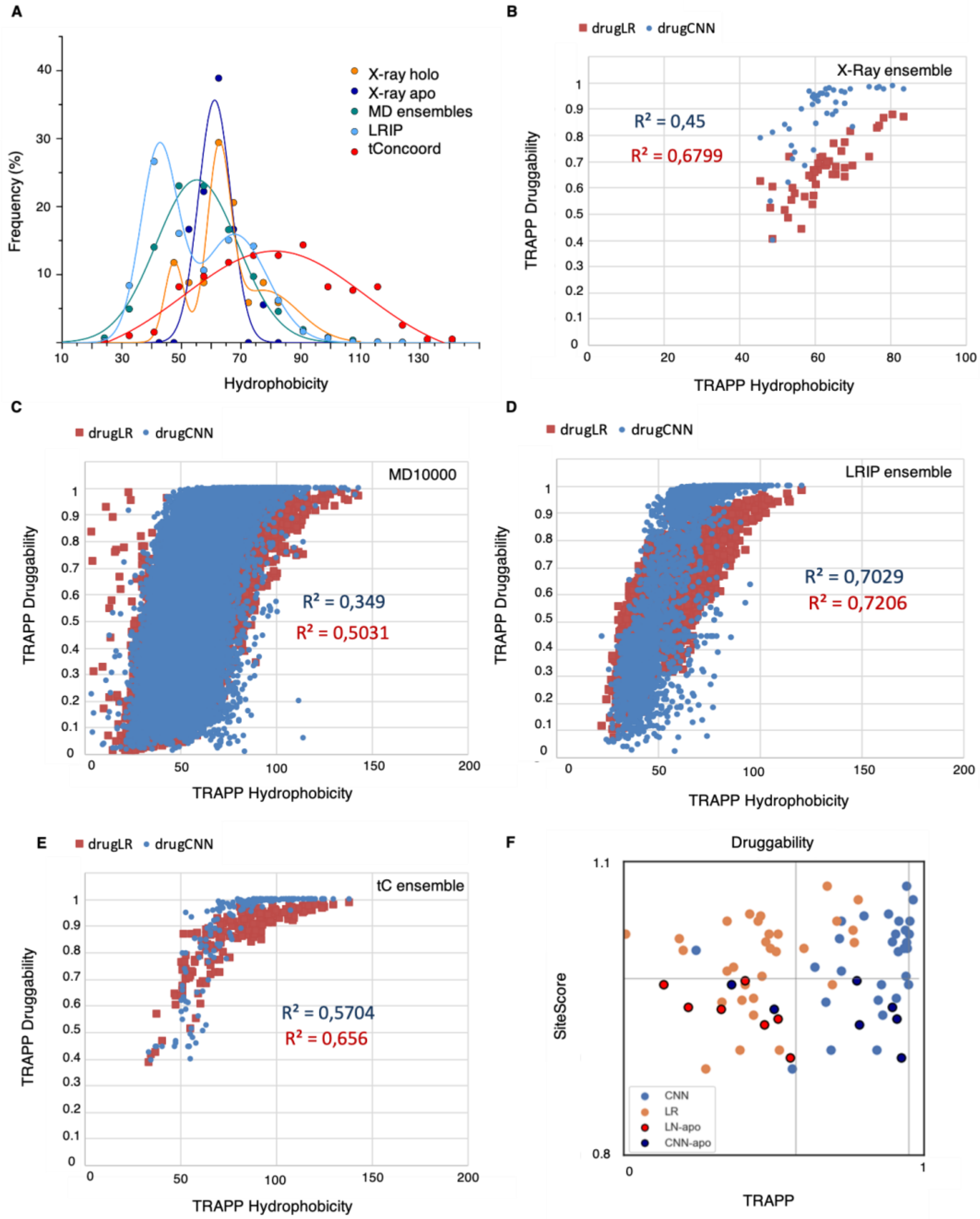


**Figure S6. Analysis of Mpro hydrophobicity.** Distribution of the hydrophobicity of the active site of different sets of Mpro (**A**) as obtained by TRAPP.[2] The data were fitted by using Gaussian functions and the Multiple Peaks Fit option available in the Origin suite. The corresponding data obtained using SiteMap for the X-ray ensemble and the ones for the MSM ensemble are reported in Table S2-S4-S5. Hydrophobicity shows a linear correlation with druggability in the X-ray crystal structure ensemble (**B**), MD10000 (**C**), LRIP (**D**) and tConcoord (**E**) ensembles. There is a large variation of the druggability index in the LR model, while the CNN druggability index better correlates with that obtained in DScore and SiteScore. From nine structures with SiteScore > 1 (PDB IDs 5REJ, 5RFS, 5RFW, 6LU7, 6M2N, 6W63, 6WNP, 7BQY, and 7BUY), six also have CNN scores above 0.95. **(F)** Comparison of the druggability predictions: SiteScore vs. TRAPP druggability scores (using LR and CNN methods) for the deposited PDB structures listed in Table S1. The *apo* structures are indicated; all others are *holo* structures.

**Table S4. Correlation analysis.** Correlation as given by $R^2$ between various binding site properties and SiteScore/DScore as calculated from SiteMap tool for X-ray crystal structures (yellow cells) and the four-state-model of the MSM ensemble (green cells). The three-state model has similar trends (data not shown).

| $R^2$ | volume | exposure | enclosure | contact | phobic | philic | balance | don/acc |
|---|---|---|---|---|---|---|---|---|
| **SiteScore** | 0.16 | 0.25 | 0.00 | 0.01 | 0.58 | 0.25 | 0.53 | 0.13 |
| **Dscore** | 0.10 | 0.12 | 0.02 | 0.00 | 0.62 | 0.41 | 0.62 | 0.13 |
| **SiteScore** | 0.29 | 0.23 | 0.12 | 0.11 | 0.04 | 0.01 | 0.02 | 0.01 |
| **Dscore** | 0.29 | 0.23 | 0.12 | 0.11 | 0.04 | 0.01 | 0.02 | 0.01 |

**Table S5. Correlation analysis.** Correlation as given by $R^2$ between various binding site properties and TRAPP LR/CNN models for X-ray crystal structures (yellow cells), the LRIP/tC ensemble (green cells), and the MD structures (blue cells). Values normalized by the binding site volume are shown in parentheses. Note that due to the grid-based method for the computation of pocket properties used in TRAPP, some correlation between pocket features and pocket volume is expected.

| $R^2$ | volume | exposure | Pos-charge | Neg-charge | h-donor | h-acceptor | Hydro phob | aromatic |
|---|---|---|---|---|---|---|---|---|
| **LR** | 0.69 | -0.19(-0.64) | 0.02(-0.01) | 0.1(-0.05) | 0.37(-0.42) | 0.38(-0.37) | 0.82(0.55) | 0.22(0.11) |
| **CNN** | 0.75 | -0.02(-0.62) | 0.02(-0.0) | 0.36(0.18) | 0.46(-0.41) | 0.61(-0.03 | 0.67(0.30) | 0.04(-0.24) |
| **LR** | 0.77 | -0.49(-0.77) | 0.27(0.22) | 0.4(-0.38) | 0.57(-0.51) | 0.65(0.11) | 0.85(0.52) | 0.72(0.29) |
| **CNN** | 0.86 | -0.39(-0.80) | 0.15(0.1) | 0.53(-0.34) | 0.74(-0.31) | 0.83(0.27) | 0.83(0.18) | 0.6(0.06) |
| **LR** | 0.49 | -0.08(-0.36) | -0.15(-0.25) | 0.06(-0.22) | 0.19(-0.56) | 0.26(-0.49) | 0.69(0.46) | 0.46(0.16) |
| **CNN** | 0.5 | -0.04(-0.37) | -0.08(-0.16) | 0.29(0.07) | 0.29(-0.38) | 0.36(-0.19) | 0.63(0.3) | 0.40(0.15) |

### SECTION S4. Virtual Screening.

In this section, cheminformatics analysis of the sample library (subsection S4.1, 4.2), structure selection (subsection 4.3), as well as, results from Glide virtual screening (subsection S4.4) along with details on the FRED virtual screening and relative PLIF (subsection S4.5) are provided. Fred and Glide EF/ROC tables are downloadable from DOI 10.5281/zenodo.4299967.

#### S4.1 The sample library diversity and active molecules chemotypes.

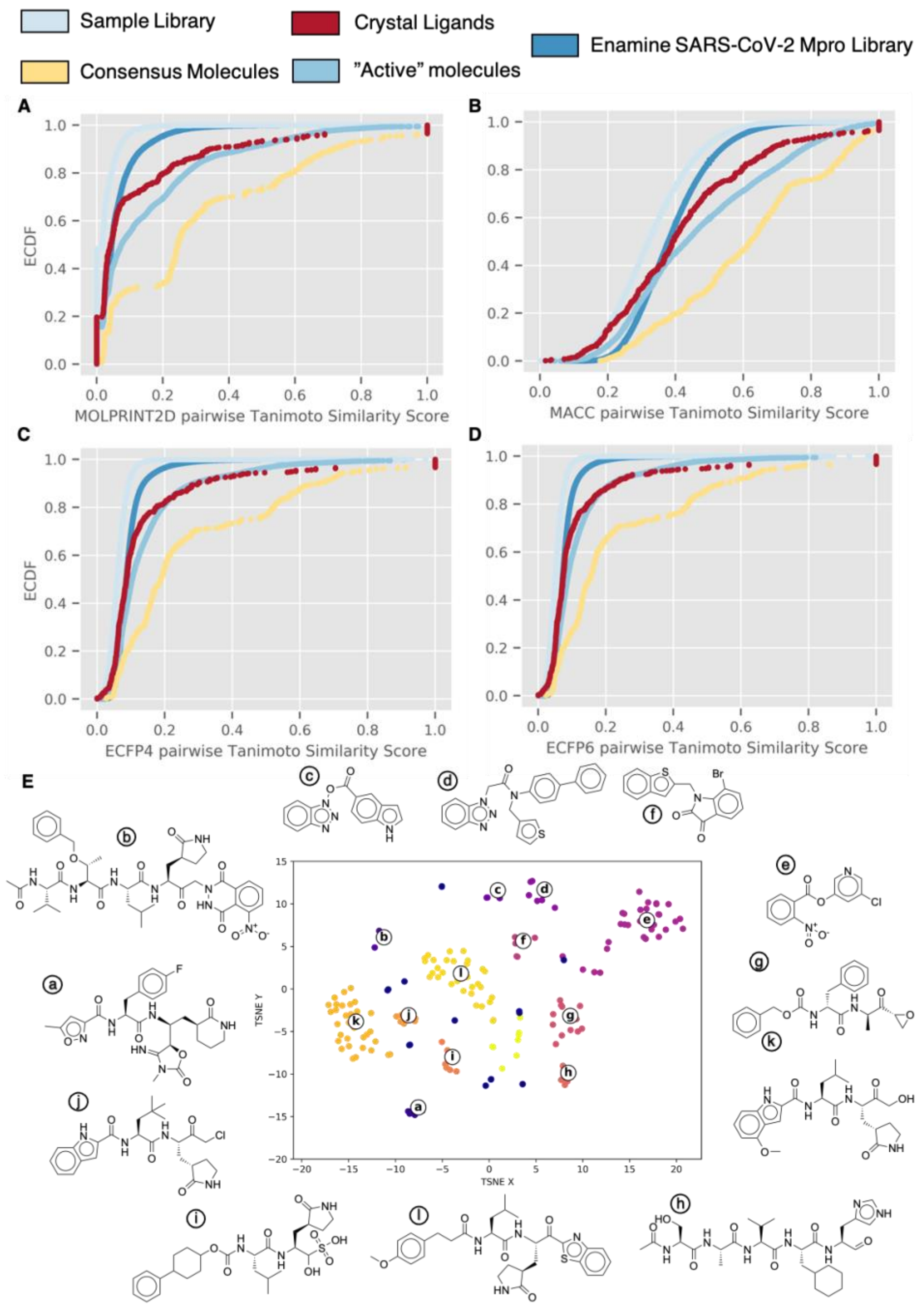


**Figure S7. Empirical Cumulative distribution function of the pairwise similarity of different ligand sets in terms of Tanimoto similarity measure.** Fingerprints used are: (**A**) molprint, (**B**) MACCS keys, (**C**) ECFP with exhaustiveness 6, and (**D**) ECFP with exhaustiveness 4. (**E**) t-Distributed Stochastic Neighbor Embedding (t-SNE) plot of the active molecules. Embedding is based on the 2048-bit Morgan fingerprint with a radius of 3. Scikit-learn t-SNE implementation with a Tanimoto distance metric was used (see supporting info Equation S1). HBDSCAN Clustering of t-SNE plot was applied resulting in 12 clusters.[3] Molecules not fitting into any cluster according to the algorithm are shown in blue and are distributed across the plot.

The diversity of the ligand library used in this work is analyzed by looking at the distribution of pairwise similarity between the molecules used in this work and the library of Enamine tailored for Mpro. We considered here our sample library, the molecule set denoted as 'active', the consensus molecules and finally the crystallographic ligands. A small variety of chemical fingerprints were considered for calculating the distance matrix using the Tanimoto coefficient as the similarity metric. The Tanimoto similarity is calculating the similarity of two fingerprints according to equation.[4] We chose to compare different types of fingerprints due to the known dependency of similarity calculations on the fingerprint type and metric used[5]. The distributions are compared by plotting the empirical cumulative distribution function (ECDF in the following paragraph, see Figure S7 A-D).

Three of the four ECDFs show a very steep rise between Tc of 0 and 0.4. The majority of similarity scores are therefore below 0.4, which highlights the dissimilarity of ligands in the databases chosen. The trends in between libraries are similar as well: the sample library is the most diverse set, followed by the Enamine. The libraries containing the crystallographic ligands, and our training set are less diverse. Looking at the ECDF generated with the MACC fingerprints, we see an apparent overestimation of the similarity between molecules. Still, the relative positions of the different distributions remain the same.

**T-SNE clustering of the molecules denoted as active in our virtual screenings**

For the ligand set denoted as active in our virtual screenings we calculated 2048-bit morgan fingerprint and used the t-SNE algorithm to plot the data set into two dimensions. Furthermore, we clustered the ligands according to the HBDSCAN clustering implemented in the scikit-learn package in an in-house python script.[3, 6] The clustering resulted in 12 clusters (see Figure S7E).

The dataset shown is dominated by peptidomimetics of different length (clusters a,b,g,k,h,l,i,j), but there are also two clusters containing diarene-esters (c,e). Clusters d,f consist of benzothiophenes and benzotriazoles.

**S4.2. Tanimoto and Dice Similarity metrics.**

$$T_c(A,B) = \frac{A \cdot B}{|A|^2 + |B|^2 - A \cdot B}$$
$$T_d = 1 - T_c$$

**Equation S1**. Tanimoto Distance and Similarity metric as implemented in t-SNE plot. A and B are bit vector representations of the chemical fingerprint of molecule A and B. Values of the Tanimoto similarity are between 0 (low) and 1 (high) similarity.

$$D_c = \frac{2|A \cdot B|}{|A|^2 + |B|^2}$$

**Equation S2**. Dice Similarity metric as used in fingerprint similarity score. A and B are bit vector representations of the chemical fingerprint of molecule A and B. Values of the Dice similarity are between 0 (low) and 1 (high) similarity.

**S4.3. Structure Selection.**

*Selection from the MD10000 snapshots and LRIP/tC ensemble***.** Structures with a druggability score above 0.9 in both CNN and LR models (3995 structures, see methods) and from MD (10,000 frames from the binding pocket of chain A) were first selected. The binding site in chain B was not used for the selection criteria since it displays significantly different behavior in MD with respect to the binding site in chain A (see Section S3 and Figure 2C of the main text). The top 10% of structures (216 structures based on site A druggability) were then clustered by the similarity of the residues lining the binding pocket as detected by TRAPP (see Figure S3C-E and Methods for more details).

Clustering showed that in some high-druggability conformations, the C-terminal tail is not involved in binding site formation (clusters 0, 2, 6; structures c, e, f; residue contribution to clusters shown in Figure S3A), while in others, it lines the pocket. The latter group generally has a larger pocket size compared to the clusters where the C-terminus is far from it (Figure S8B, Figure S9D). From each of the 8 clusters generated, one representative was chosen.

As one can see from the heatmap of cluster composition and pocket property visualization for each cluster (Figure S8B), clusters 1, 3-5, and 7 comprise structures with C terminus of chain B lining the binding pocket, which efficiently increases the pocket volume. Clusters 0, 2, and 6 differ by contribution of residues from ASL1 and ASL2. To select one representative from each cluster for docking, the conformations with summed LR and CNN druggability scores greater than the 95% quantile from each of the clusters were inspected manually. Conformations with close similarity to others and those with contributions to druggability from residues distant from the pocket were discarded. For cluster 5, the one with the lowest druggability in the top 10% scoring conformations, the second highest-scoring conformation was chosen.

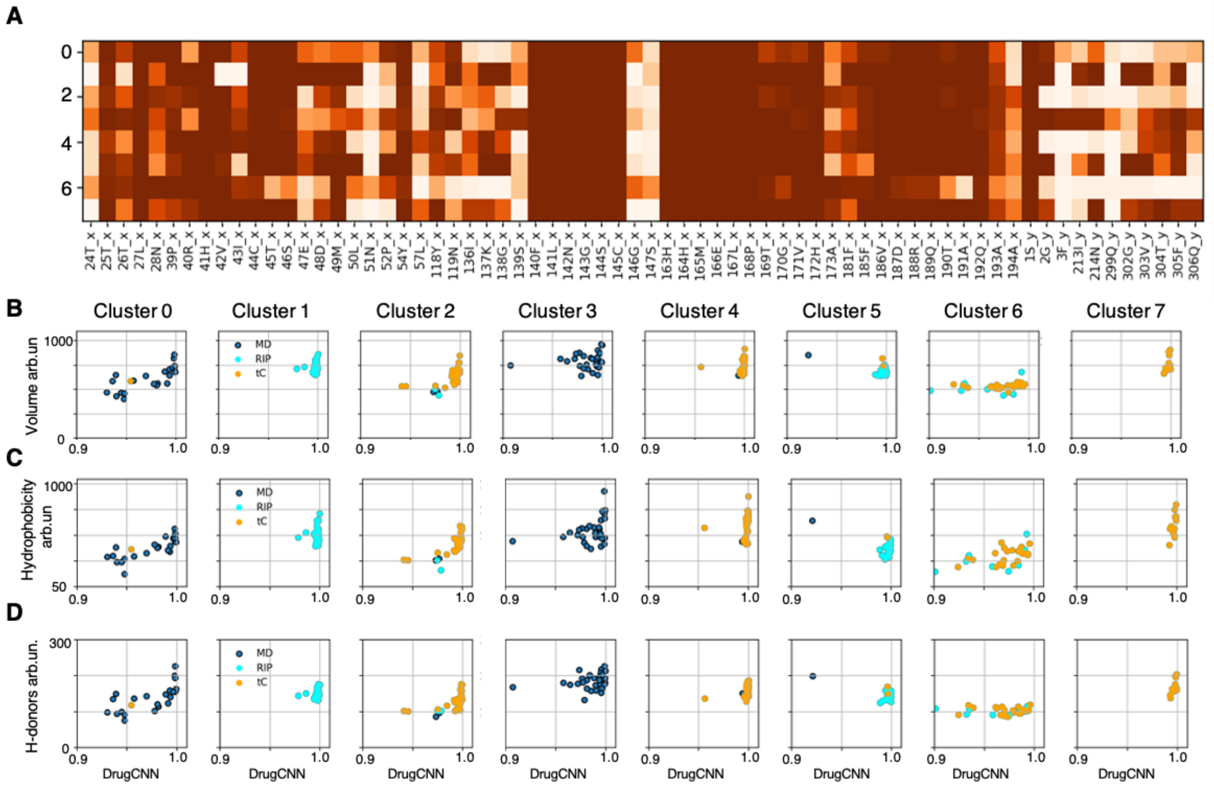


**Figure S8. Results of clustering of the top 10% of druggable structures (according to the TRAPP CNN and LR scores) from 10,000 frames (i.e. every 1 ns) of the 100 µs-long MD and an ensemble of structures generated using, tConcoord and LRIP methods.** (**A**) Contributions of the binding site residues averaged over the cluster members is shown for eight clusters by color varying from white (no contribution) to dark orange (contribution from all cluster members); (**B-D**) Distribution of three pocket properties vs. CNN druggability score for each of the eight clusters.

Three regions in the pocket appear to contribute strongly to the druggability score of the CNN model in many of the selected structures.

1) The pocket wall formed by ASL 1&2 opens up with unfolding of ASL1 and results in a positive influence on druggability (see Figure S9E; structures c, d, e, f, and h). Binding of this region in the closed form appears in some experimental X-ray crystal structures (PDB IDs 5R80 and 5RG1).
2) Opening of ASL2 and P168 allows access to a hydrophobic pocket region (see Figure S9), increasing druggability. This motion is observed in structures b, c, d, f, g, and h (see Figure S9E) and appears in tConcoord and MD structures. In MD simulations, extension of this motion and formation of a helix occur additionally (see Figure S4).
3) Contributions of the C-terminal tail are present in the structures selected in two forms: either by reaching across the pocket, contributing to the druggability of the pocket center (structure b, of note: this happens in most structures of cluster 3) or by lining the pocket in various places (structures a, d, and g).

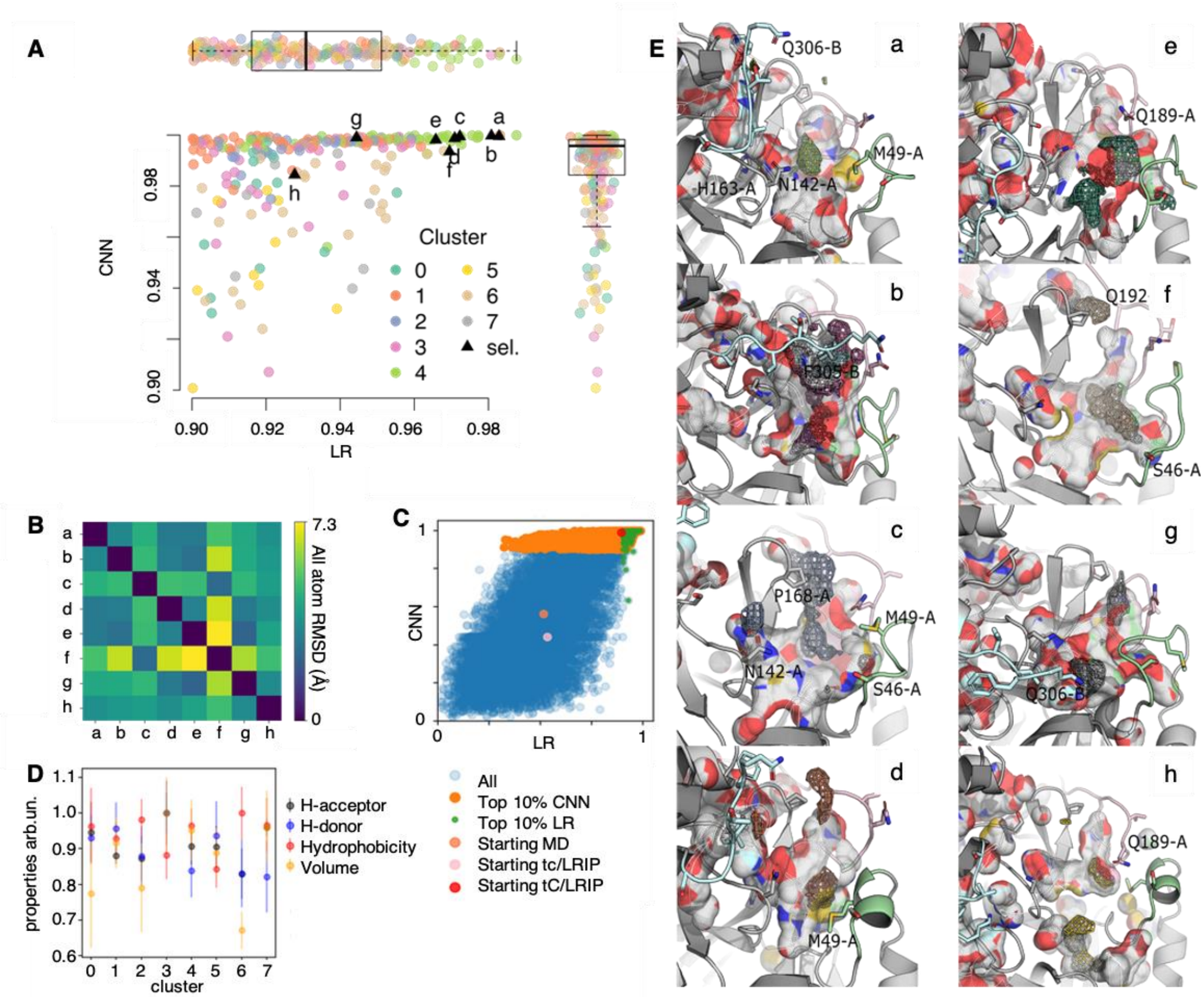


**Figure S9. Selection of the structures for docking from the set of MD frames and LRIP/tC ensemble.** (**A**) distribution of druggability scores plotted for highly druggable structures colored by cluster; a-h points denote structures selected from clusters 0-7, respectively, as given in Table S6; (**B**) heatmap of pocket RMSD between the selected cluster representatives calculated with pymol; (**C**) TRAPP druggability score distribution with the CNN model plotted vs the LR) of all computationally generated structures; (**D**) pocket properties normalized to the pocket size and averaged over each cluster with the standard deviation shown by bars; (**E**) the representatives (labeled a to h) are shown as cartoons with the pocket surfaces shown and colored by atom type. The side chains of relevant residues are shown as licorice and the region contributing most (30%) to the CNN druggability score is shown by iso-mesh, colored according to the cluster number as in panel A. ECL1 is colored light green, ECL2 light pink and the C-terminal tail cyan.

### S4.4: Glide virtual screening

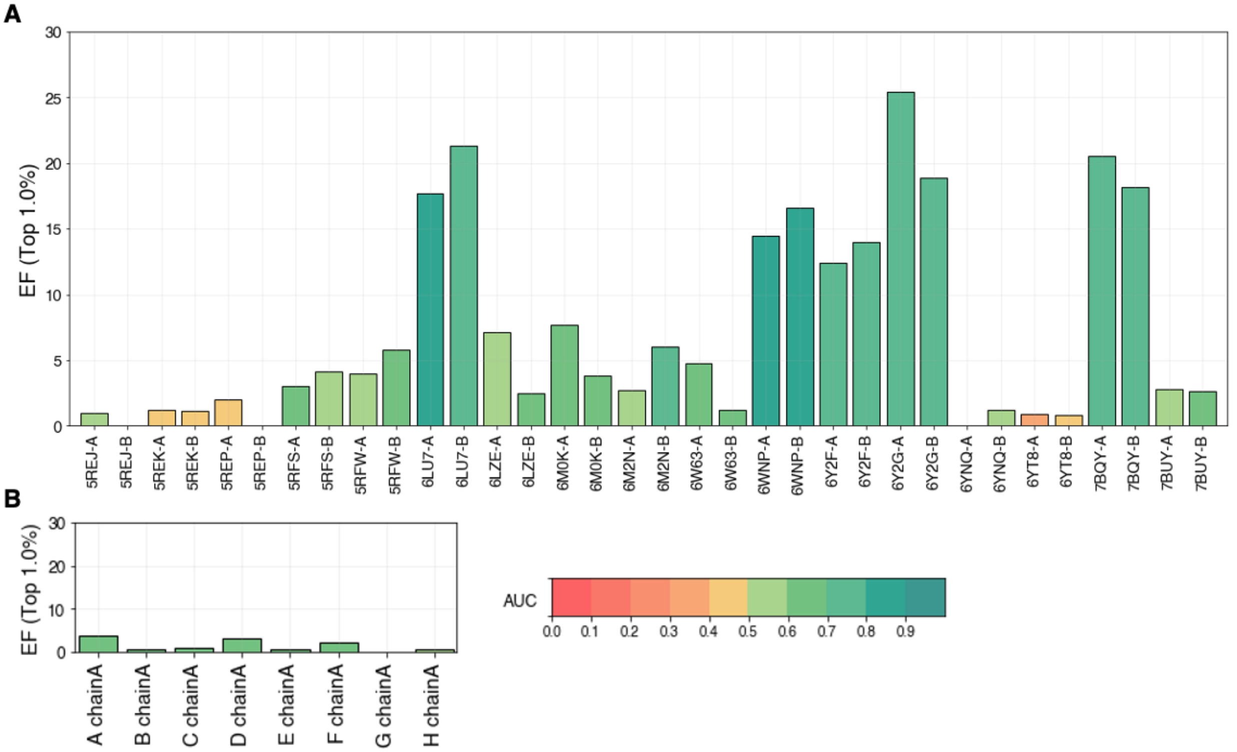


**Figure S10. Glide docking results.** EF for the "active" molecules found in the best 1% of the results of the GLIDE virtual screenings performed on each chain found in the crystal structures (**A**), in the "DrugScore selection" group (Section S3.4) (**B**) and in the MSM selection, i.e. 10 structures with druggability index > than 0.9, extracted from the four-state model MSM ensemble.

### S4.5. Fred virtual screening.

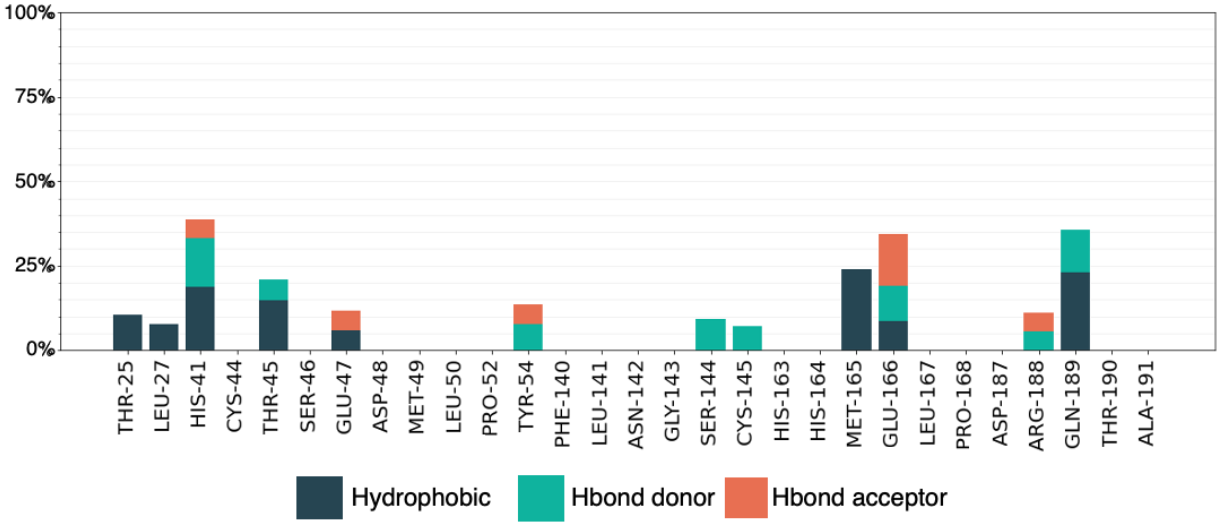


**Figure S11. PLIF of the top 1% molecules docked on selected structures from the four-state MSM ensemble.**

**Table S6. Feature space data and druggability scores calculated with TRAPP for the representative structures selected in figure 4.** Three druggability scores are given, the linear regression (LR) score, the convolutional neural network (CNN) score and the sum of both. The summed score was used for ranking of the structures.

| Structure | a | b | c | d | e | f | g | h |
|---|---|---|---|---|---|---|---|---|
| Cluster | 4 | 3 | 2 | 1 | 0 | 6 | 7 | 5 |
| drugSUM | 1.982 | 1.980 | 1.971 | 1.969 | 1.963 | 1.963 | 1.943 | 1.912 |
| drugLR | 0.982 | 0.980 | 0.972 | 0.970 | 0.965 | 0.969 | 0.944 | 0.927 |
| drugCNN | 0.999 | 0.999 | 0.999 | 0.998 | 0.998 | 0.993 | 0.999 | 0.984 |
| Volume | 806.2 | 955.2 | 698.2 | 841.5 | 706.8 | 533.6 | 681.5 | 452.8 |
| Exposure | 14.94 | 13.37 | 14.62 | 14.16 | 9.42 | 12.70 | 9.15 | 12.92 |
| pos-charge | 1.82 | 0.60 | 1.37 | 11.30 | 0.19 | 0.04 | 0.16 | 0.00 |
| neg-charge | 26.33 | 49.86 | 15.00 | 30.89 | 27.20 | 18.64 | 37.86 | 12.47 |
| h-donor | 135.8 | 175.0 | 131.3 | 153.9 | 135.3 | 80.7 | 139.7 | 75.0 |
| h-acceptor | 161.6 | 213.3 | 134.3 | 176.7 | 155.7 | 103.7 | 163.4 | 97.6 |
| Hydrophob | 118.9 | 124.6 | 106.8 | 113.8 | 94.1 | 81.2 | 93.6 | 71.1 |
| Aromatic | 29.75 | 52.53 | 23.85 | 24.27 | 13.67 | 17.08 | 11.09 | 19.33 |
| Metal | 0 | 0 | 0 | 0 | 0 | 0 | 0 | 0 |

**SECTION S5. Binding site hot-spots distribution across MD: comparison with X-ray crystal structures and impact on the EF**

Detailed analysis on the distribution and compactness of the interaction hotspots of the MD conformations and the impact on the EF is here offered. In particular we elucidate if: binding site conformations similar to the well-performing structures are sampled during MD (i) and if these perform similarly to the well-performing structures (ii). We found that binding site conformations similar to the well-performing ones also occur in the MD. Indeed, the Root Mean Square Deviations (RMSD see Equation S3) with respect to the well-performing structures (structures as defined in Section 2 of the main text) can be lower than 1.5 Å for the binding site residues (residues which occurred more than 55% in Figure 2D in the main text). However, these occur with a very low frequency. Despite the low RMSD, the shape of these binding sites substantially differs from those of the well-performing structures (Figure S13) and they do not perform as well as the well-performing structures.

$$\mathrm{RMSD}(v,w) = \sqrt{\frac{1}{n}\sum_{i=1}^{n}((v_{ix}-w_{ix})^2+(v_{iy}-w_{iy})^2+(v_{iz}-w_{iz})^2)}$$

**Equation S3.** Root Mean Square Deviations with $v$ and $w$ indicating atom coordinate vector of the same atom in two different MD frames.

We obtained similar results when trying to find MD snapshots with a pocket similar to that of PDB ID 6LU7, using the number of atoms of each binding site residue lining the binding cavity (as computed by TRAPP): the similarity between the crystal structures was found to be higher or comparable with the most similar MD snapshots. Accordingly, in the screening assessment no high-enrichment structures were found from the selected list (DOI 10.5281/zenodo.4299967).

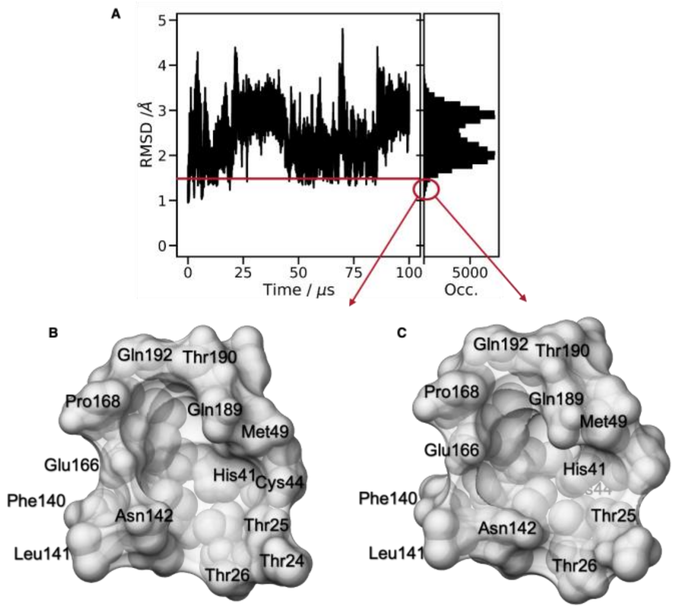


**Figure S12. Analysis of the D. E. Shaw MD trajectory.** (**A**) Root Mean Square Deviation (RMSD) overall the D. E. Shaw MD trajectory with respect to the average of the X-Ray structures (left) and (**B**) corresponding distribution. The red line and circles in left and right plot, respectively, indicate the cut-off of 1.5 A. (**C**)(**D**) MD frames with binding site conformations similar to the well-performing structures.

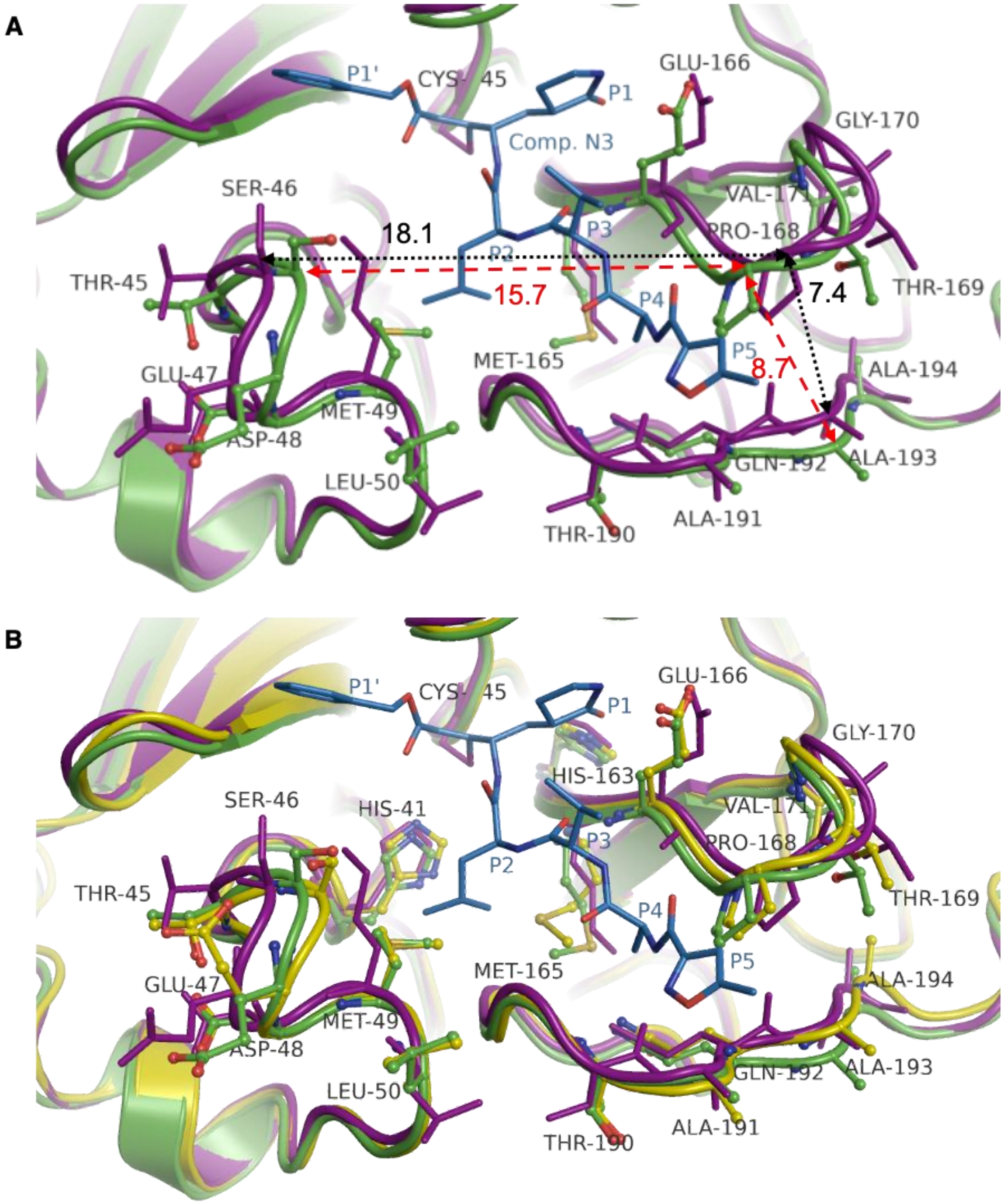


**Figure S13. Structural Differences between well-/poorly- performing structures.** (**A**) Superposition of the crystal structure of SARS-CoV-2 Mpro *apo*, PDB ID 6WQF: green carbons, *holo*, PDB ID 6LU7: purple carbons. (**B**) Superposition of well-/poorly- performing and *apo* structures. This figure was based on the scheme published in ref. Kneller et al..[7] We only show here a single structure for each category (i.e. well-, poorly- performing and *apo*). Similar results were obtained by comparing other structures (data not shown).

### SECTION S6. Chemical Space.

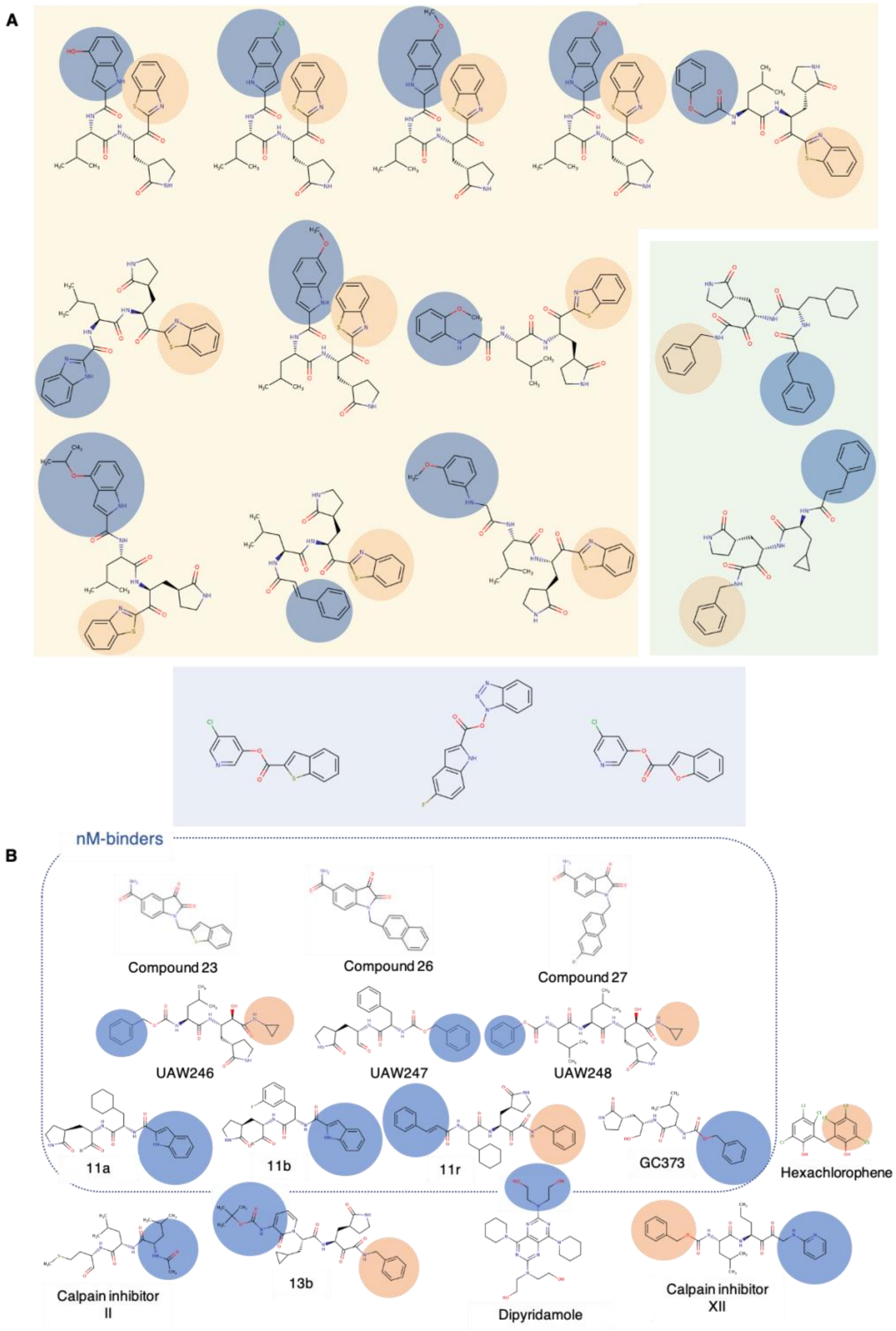


**Figure S14. Analysis of the chemical space.** (**A**) “Consensus active” molecules and (**B**) sub-µM and nM inhibitors of SARS-CoV-2 Mpro with a binding pose. Compounds 23, 26, and 27 were obtained from literature[8] during the submission process. Molecules in panel (**A**) are divided into three clusters as described in Section S4. The blue circles highlight the moieties buried in the cavity defined by residues Glu166, Pro168, Gln182, Thr190 and Gln189, while the orange circles show the portion of the molecules in proximity to Thr24, Thr25, Thr26.

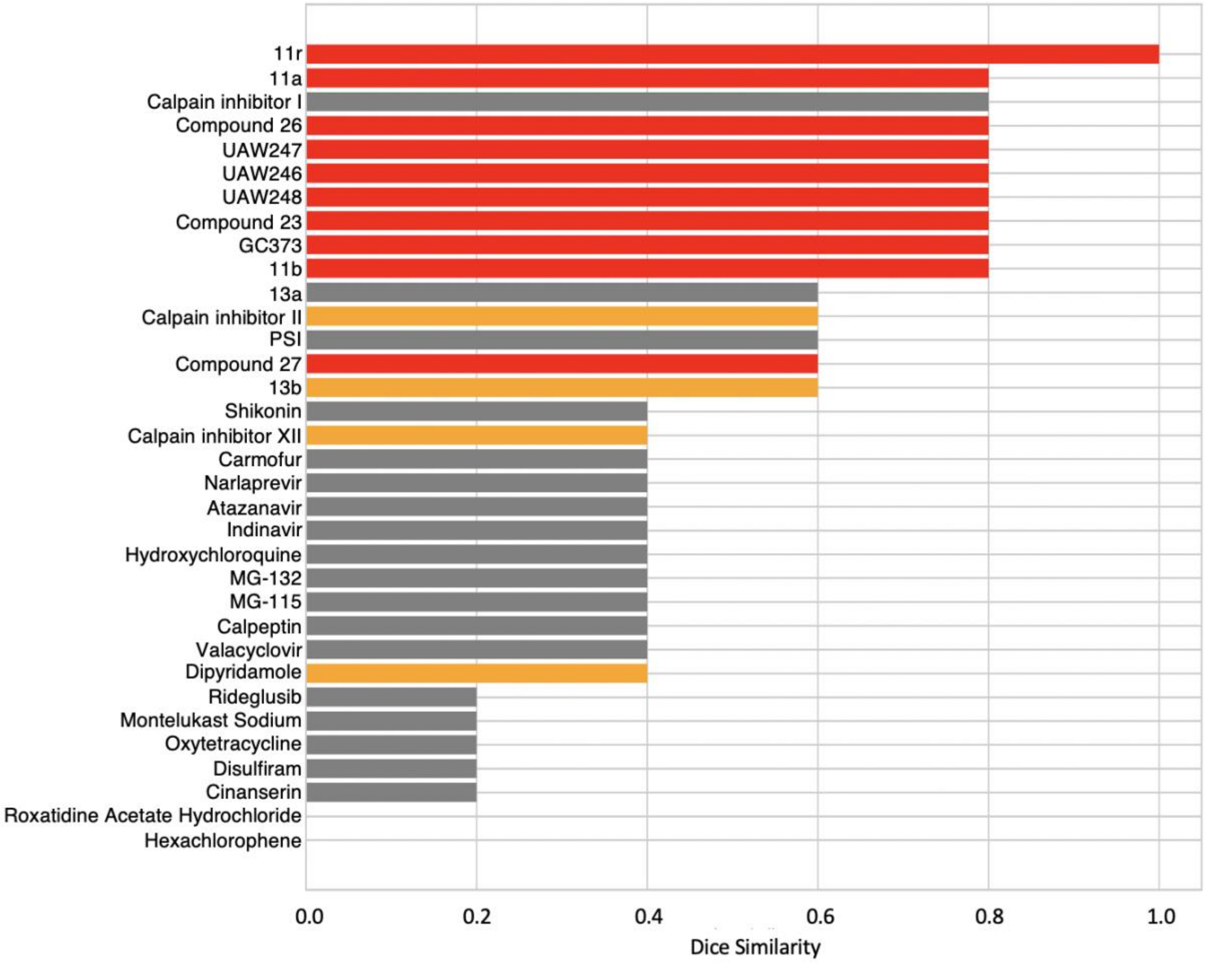


**Figure S15. Bar plot of Dice similarity index between the active-pharmacophore and the docked pose of known SARS-CoV-2 Mpro inhibitors.** The high nanomolar range (400 nM < IC50 ≤ 1000 nM) inhibitors are shown in orange, while ligands with IC50 ≤ 400 nM are shown in red. Micromolar inhibitors discovered in literature[8, 9] after 20$^{th}$ October 2020 were not included.

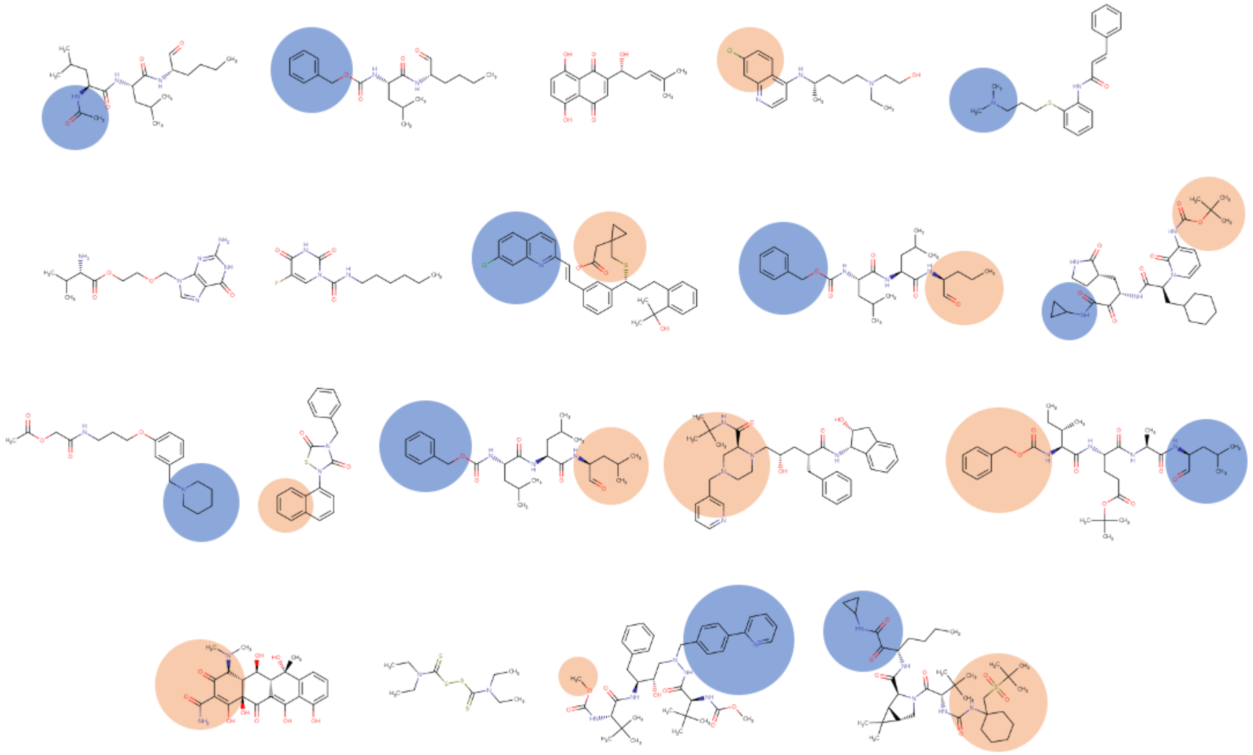

**Figure S16. Molecules in the micromolar inhibitor group.** The blue circles highlight the moieties buried in the cavity defined by residues Glu166, Pro168, Gln182, Thr190, and Gln189, while the orange circles show the portion of the molecules in proximity to Thr24, Thr25, and Thr26.

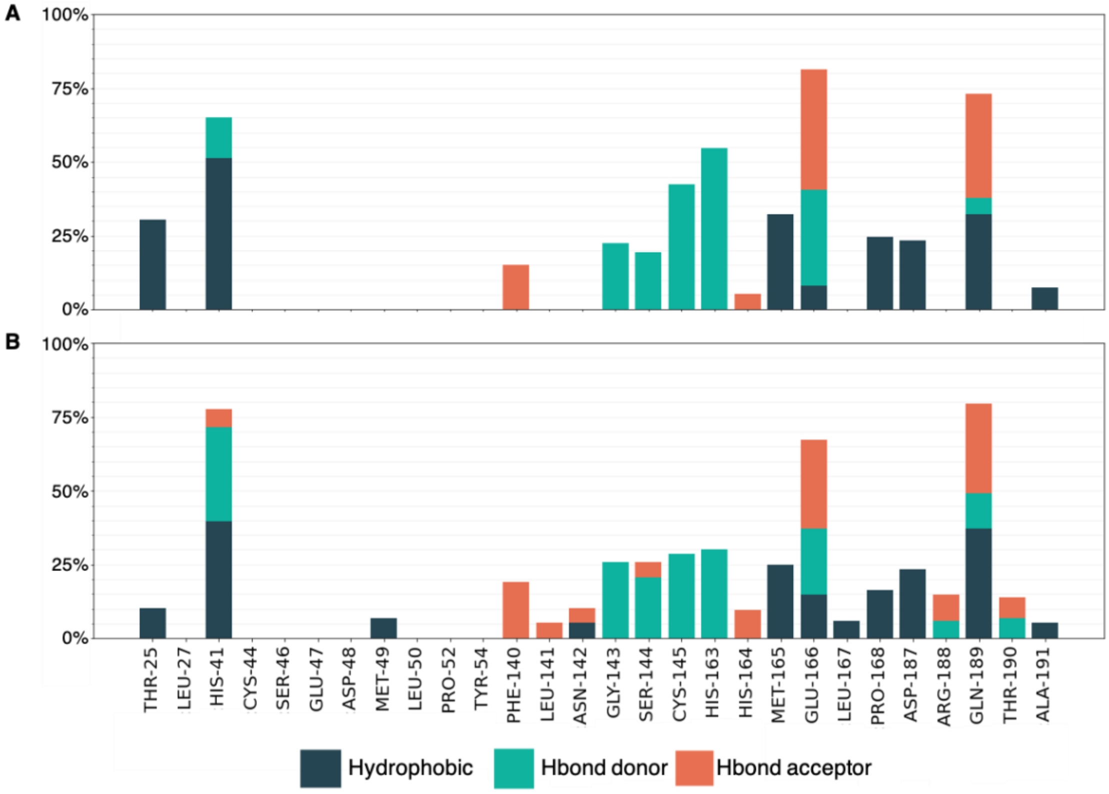


**Figure S17. PLIF analysis.** (**A**) PLIF of the top 1% consensus molecules across the well-performing structures, (**B**) PLIF of SARS-CoV-2 Mpro inhibitors in the sub-micromolar range (400 nM < IC50 ≤ 1000 nM)

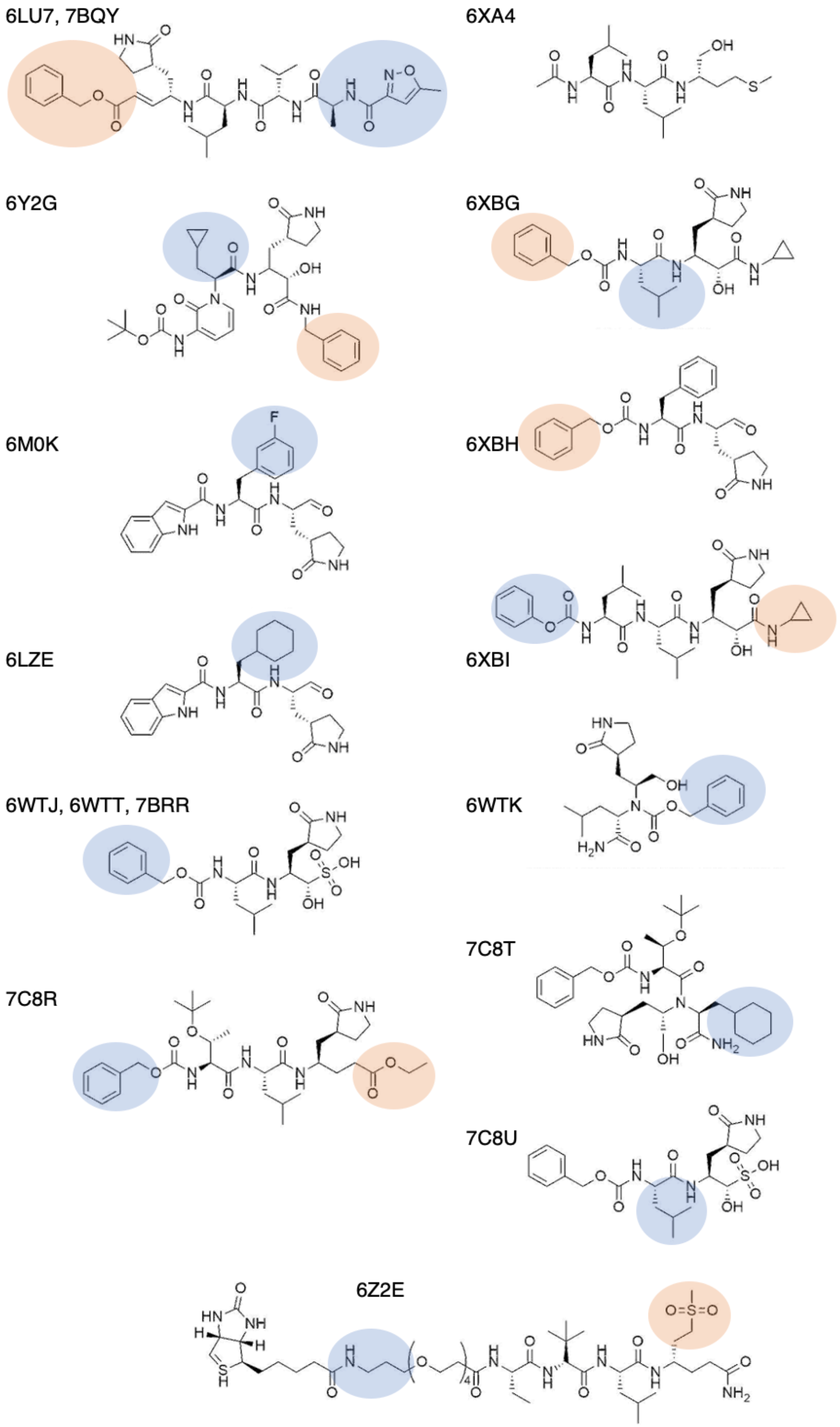


**Figure S18. Co-crystalized ligand structures and their corresponding PDB ID.** The blue circles highlight the moieties buried in the cavity defined by residues Glu166, Pro168, Gln182, Thr190 and Gln189, while the orange circles show the portion of the molecules in proximity to Thr24, Thr25, and Thr26.

### SECTION S7. Molecular Dynamics simulation on PDB ID 7BRR.

MD simulations were performed with the Desmond version implemented in the Schrödinger package. The preparation of the protein-ligand complex was accomplished with the Protein preparation wizard with standard parameters. The complex was first immersed into a TIP3P water box, extending 10 Å beyond any of the complex's atoms in each direction. Counter ions (6 $Na^+$ Ions) were added to neutralize charges. The salt concentration was set to 0.15 M sodium and chloride ions to approximate physiological condition. After the standard 5 step relaxation protocol, the MD was performed in the NPT ensemble at a temperature of 300 K and 1.01 bar pressure over 500 ns with recording intervals of 100 ps, resulting in 5000 frames.[10] The OPLS-3e force field was used.[11]

During our docking experiments, we found a difference between the pose of compound 11r in the crystal structure and our docking pose. In the crystallographic pose, the phenyl moiety is arranged in such a way that it faces the solvent and appears not to have a single contact with the protein residues despite being hydrophobic. We wanted to see if the placement of the phenyl might change during the course of a short MD calculation.

Indeed, the phenyl moiety of compound 11r appears to rearrange during the 500ns MD trajectory performed. In the majority of frames, the phenyl moiety is in close contact to a subpocket defined by the two loops defined by residues Glu166, Pro168, Thr190, Ala191 and Gln192. Furthermore, the dihedral angle values between atoms C-O-O-C (marked in red color) is highly populated around the angle -180 and 180, which is a similar value to the one identified by our docking calculation.

The high mobility of the phenyl moiety is also reflected in the RMSF plot of the ligand, which reaches highest values on atom numbers 6 to 9. During the course of the MD simulation of 500 ns, the RMSD of the protein heavy atoms reaches an equilibrium around the value of 3 Å and the ligand atom's position around 2.2 Å.

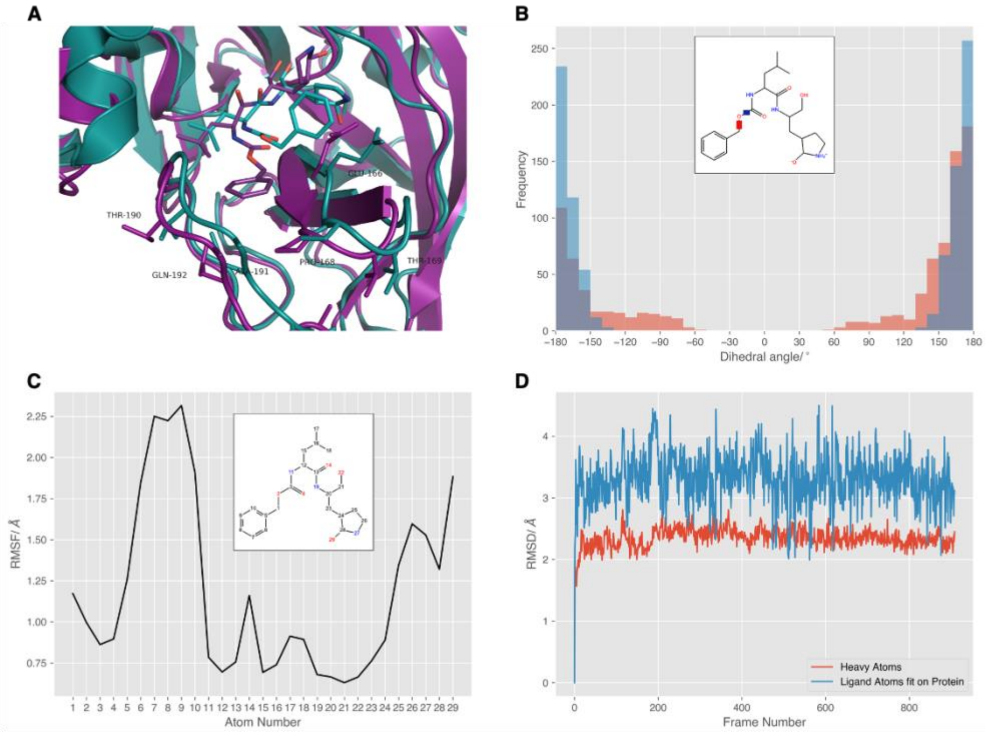


**Figure S19. Analysis of the trajectory.** (**A**) The phenyl moiety of compound 11r shows rearrangement during 500ns MD trajectory (MD structure shown in purple, initial crystal structure frame's atoms are shown in cyan). It fills a subpocket defined by the residues Glu166, Pro168, Thr190, Ala191 and Gln192. (**B**)(**C**) During the MD, the dihedral angle between the atoms C-O-O-C (marked in red) and N-(CO)-O-C (marked in blue) are highly populated around the angle -180 and 180, which fits the representative MD orientation shown in panel **A** (purple atoms). The high mobility of the phenyl moiety is also reflected in the RMSF plot of the ligand, which reaches highest values on atom numbers 6 to 9. (**D**) During the MD simulation of 500 ns the RMSD of the protein and ligand heavy atoms reaches an equilibrium around the value of 3 and 2.2 Å respectively.

### SECTION S8. Excluded molecules from the consensus.

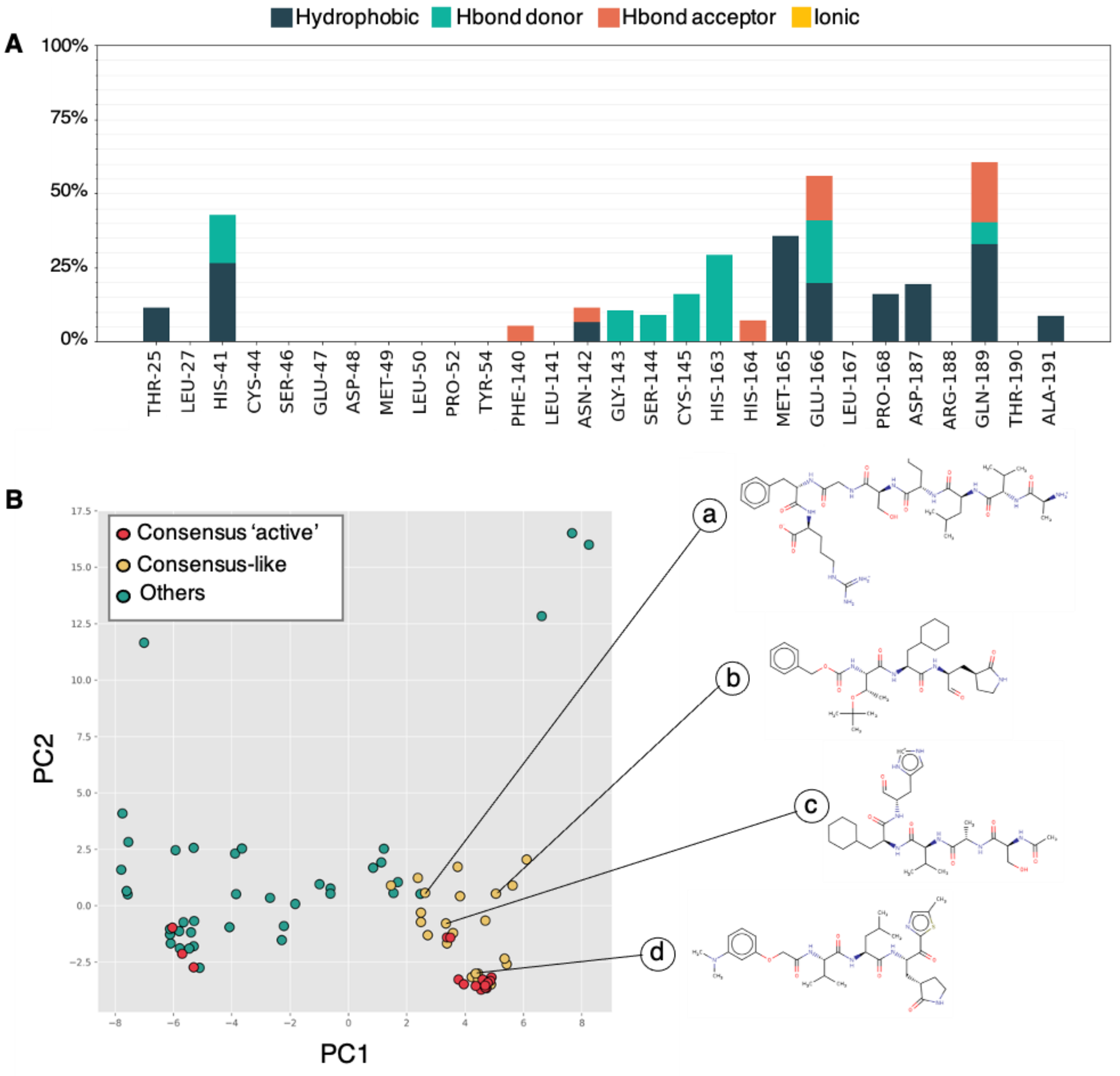


**Figure S20. Molecule analysis.** (**A**) PLIF of the "active" molecules not in the consensus docked on the well-performing structures. (**B**) Principal Component Analysis (PCA) of the MACCS fingerprint generated for the "active" molecules in the sample library. In red, the molecules in the consensus 'active' group. In yellow the "active" molecules the co-clusterize with the molecule in the consensus. In green, all the other "active" compounds.

### SECTION S9 Nanomolar inhibitors identification.

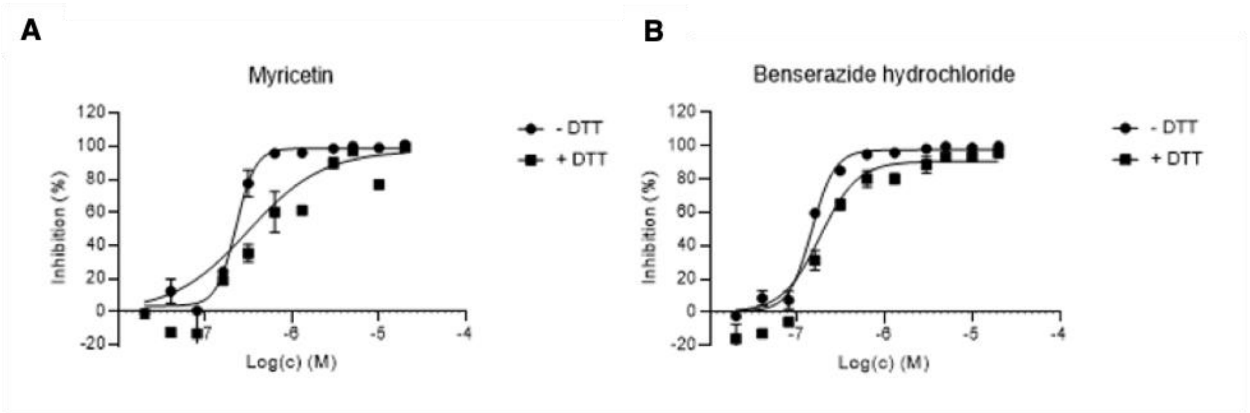


**Figure S21. Dose dependent effect on Mpro activity.** Dose dependent effect of Myricetin (**A**) and Benserazide hydrochloride (**B**) on SARS-CoV-2 Mpro activity.

#### S9.1 Induced Fit Docking Procedure Myrecitin.

The induced fit docking was started with the docking pose of Myricetin docked by with FRED openeye software described in the Methods section of the main text. The standard maestro protocol [12] [13] [14] [15] was applied with a cutoff for the sidechain trim of 5.0 Å centered upon the ligand. A maximum of three trimmed sidechains with the B-factor method at a cutoff of 40 $Å^2$ was chosen, which resulted in the trim of residues 49,189,44. For the docking procedure an inner grid of 10 Å was used and the docking was constrained on the ligand's heavy atoms with a restriction to 1 Å to the initial pose resulting from the FRED procedure. We chose the best scoring pose according to the IFDScore for the comparison in Figure 6C.

#### S9.2 Baicalein docking

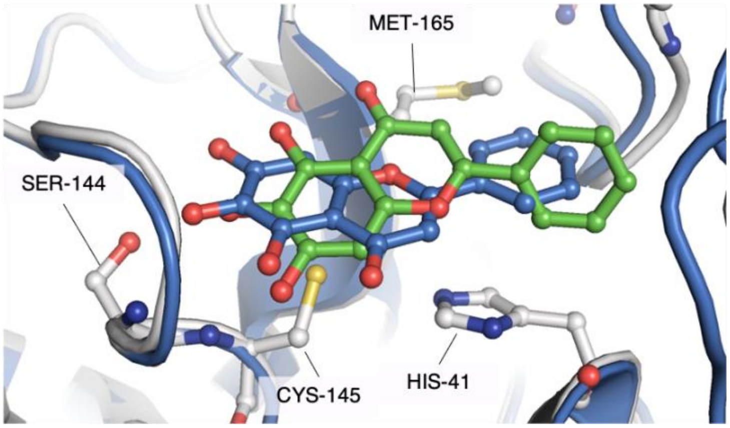


**Figure S22. Baicalein redocking.** In blue, pose of Baicalein docked on one of the well-performing receptors (6LU7, blue cartoon representation). In green, the ligand pose of Baicalein as observed in the X-ray structure with PDB ID 6M2N (protein in light grey cartoon and licorice representation). The RMSD between the docking- and the crystal structure pose is 2.93 Å. In both cases, the bicyclic ring of the molecule is in a different orientation than the one appearing in Myricetin docking- and crystal structure (Figure 6C).
